# AlphaFold reveals but sometimes distorts an organizational principle of protein folding

**DOI:** 10.64898/2025.12.30.697132

**Authors:** Madeleine F. Clore, Joseph F. Thole, Suchetan Dontha, Pramesh Sharma, Naomi Greenberg, Marie-Paule Strub, Mary Starich, Carolyn Ott, Davin Jensen, Brian F. Volkman, Matthew Coudron, Lauren L. Porter

## Abstract

Proteins can adopt many distinct conformations, yet the sequence determinants governing which structural states are accessible remain poorly understood. Using fold-switching proteins and AlphaFold as an analytical lens, we identify an organizational principle in which access to alternative structural states–and in some cases the folded state–is governed by surprisingly few amino acids, termed gating residues. Gating generalizes to single-fold proteins, indicating that conformational accessibility is often hierarchically organized around a small number of disproportionately influential residues. AlphaFold has implicitly learned this principle but amplifies and sometimes misapplies it, concentrating conformational control onto too few or incorrect residues. Guided by this principle, targeted MSA editing recovered a conformation AlphaFold confidently mispredicted, suggesting a path toward more accurate prediction of alternative conformations and mutational effects.

## Introduction

How a single protein’s conformation is specified among many possible states remains a fundamental unsolved problem in structural biology. Conformational selection underlies allostery (*1*), signal transduction (*2*), and functional diversification of protein families (*3*), yet the sequence determinants governing which structural states are accessible remain poorly understood. A central challenge is determining which amino acid residues govern conformational accessibility and whether they reflect a general organizational principle. Identifying these determinants could reveal fundamental principles of protein folding while improving prediction of conformational ensembles and mutational effects, a persistent challenge for AI-based models (*4–10*).

AlphaFold (*11, 12*) offers an unexpected opportunity to address this question. Although trained to output a single predicted structure, it must implicitly represent information governing conformational accessibility to select among alternative structural possibilities. AlphaFold combines information from the amino acid sequence with a multiple sequence alignment (MSA), allowing its internal representations to integrate both sequence-specific and evolutionary constraints. Most analyses presented here isolate single-sequence predictions, showing that the principles we identify are not driven by explicit coevolutionary information. We later demonstrate that these principles extend to MSA-based predictions.

While remarkably successful at predicting single structures and oligomeric assemblies from amino acid sequences (*11–13*), AlphaFold remains unreliable at selecting among alternative conformational states (*8, 14–17*). Rather than viewing these failures solely as limitations, we asked whether they reveal biologically meaningful principles governing conformational accessibility. As a striking example, AlphaFold confidently predicts that pro-interleukin-18 (pro-IL-18) adopts its mature conformation despite experimental evidence that the two structures differ (**Fig. 1A**) (*18*). The pro- and mature forms differ by a 36-residue N-terminal insertion (that is well-represented in the input MSA, **Fig. S1**) and share 81% sequence identity.

**Figure 1.**
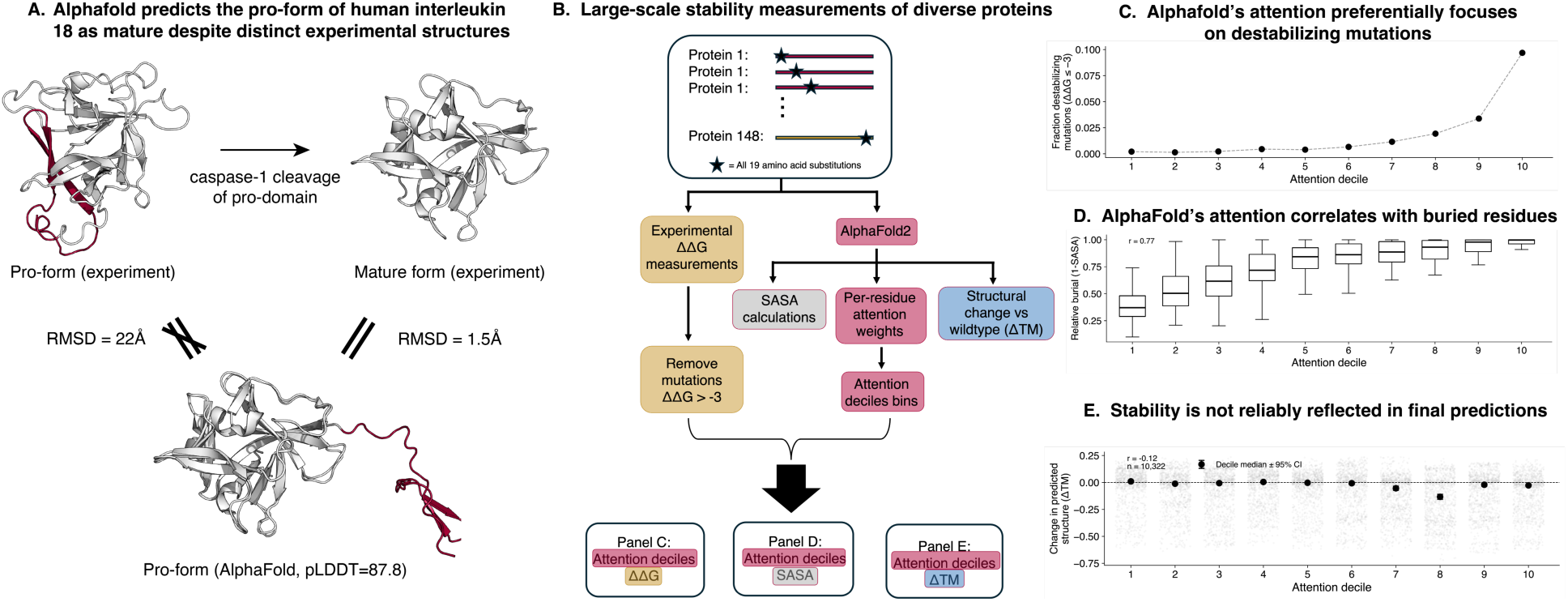
AlphaFold2 encodes information related to thermodynamic importance but does not typically propagate it to structural predictions. **(A)** AlphaFold2 incorrectly predicts the pro-form of human interleukin-18 (IL-18) in the mature conformation despite experimentally distinct structures. Cleavage of the N-terminal pro-domain produces the mature structure (top), whereas AlphaFold2 predicts the mature conformation directly from the pro-form sequence with high confidence (bottom; pLDDT = 87.8). Pro-form PDB ID: 8URV; Mature: 3WO2. **(B)** Overview of large-scale mutational analysis. Experimentally measured thermodynamic stabilities (ΔΔG) for up to 19 amino acid substitutions across 148 proteins were compared with AlphaFold2-derived quantities, including per-residue attention weights, residue burial (SASA), and predicted structural changes relative to wild type (ΔTM). **(C)** AlphaFold attention preferentially focuses on residues associated with strongly destabilizing mutations. Residues binned by attention decile exhibit monotonic enrichment of mutations with ΔΔG ≤ −3 kcal/mol, indicating that AlphaFold allocates greater attention to positions with greater thermodynamic importance. **(D)** Attention correlates strongly with residue burial across diverse proteins with destabilizing mutations (ΔΔG ≤ −3 kcal/mol, r = 0.77), suggesting that AlphaFold preferentially emphasizes structurally buried positions. **(E)** Thermodynamic information encoded within AlphaFold’s internal representations is not faithfully reflected in its structural outputs. Predicted structural changes (ΔTM) show little relationship with attention decile (r = −0.12; n = 10,322), indicating that residues receiving the greatest attention are not more likely to produce appreciable structural changes or decreased confidence (**Figure S2**) when mutated. Decile medians ± 95% confidence intervals are shown.

AlphaFold predictions—including its failures—therefore provide a window into how conformational accessibility may be represented within protein sequences. Fold-switching (metamorphic) proteins (*19–21*) provide a stringent test of conformational selection because they adopt multiple disparate structures that can be compared directly against AlphaFold predictions (*9*), and several have mutations that cause measurable shifts in conformational equilibria (*22–24*). These systems allow us to ask how AlphaFold allocates conformational control across residues and whether the positions it emphasizes correspond to biophysical determinants of conformational accessibility.

We first asked whether AlphaFold’s internal representations encode information related to experimentally measured mutational effects (Fig. 1). We focused on AlphaFold’s attention weights, which quantify the relative importance assigned to residues during structure prediction. We then analyzed these representations in experimentally characterized fold-switching proteins to determine how AlphaFold selects among alternative structural states, identifying a small subset of gating residues that hierarchically control conformational accessibility (Figs. 2–3). Next, we tested whether these predicted control points correspond to biologically meaningful determinants of conformational selection using the fold-switching proteins KaiB and RfaH (Fig. 4). Finally, we asked whether AlphaFold assigns these residues the same influence as single-folding proteins and whether understanding this bias could be used to improve incorrect conformational predictions (Fig. 5).

**Figure 2.**
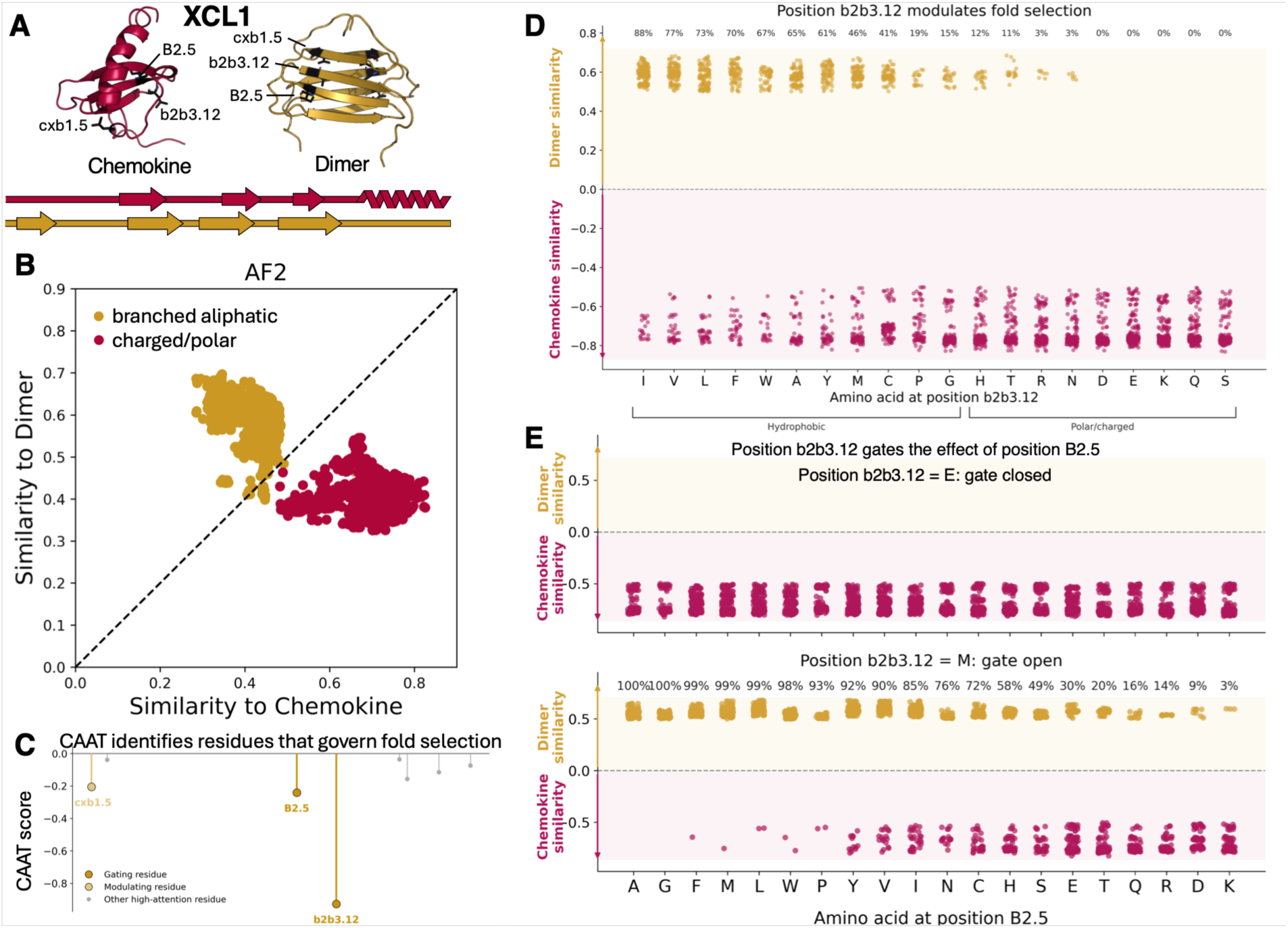
AlphaFold filters access to conformational states through sparse, hierarchically organized gating residues. (A) XCL1 adopts two distinct conformations—a monomeric chemokine fold (red, PDB ID: 1J9O) and a β- sheet dimer (gold, PDB ID: 2JP1)—with positions cxb1.5, B2.5, and b2b3.12 in black. These positions form an interaction network unique to the dimer fold. (B) AlphaFold (AF2) predictions across sequence variants separate into two discrete structural states. Variants containing branched aliphatic residues at positions cxb1.5, B2.5, and b2b3.12 are predicted to adopt the dimer conformation, whereas charged and polar residues at these positions favor the chemokine fold; this result also holds for AlphaFold3 (**Figure S4**). (C) CAAT identifies positions cxb1.5, B2.5, and b2b3.12 as the primary conformational access points governing fold selection, based on comparison of attention patterns across sequence variants. (D) Systematic amino acid substitution at position b2b3.12 reveals single-residue gating. Hydrophobic residues favor the dimer conformation (gold), whereas polar and charged residues favor the chemokine fold (red). The fraction of dimer predictions (top) ranges from 0% (D, E, K, Q, S) to 88% (I). (E) Hierarchical gating by position b2b3.12. When b2b3.12 is fixed to a chemokine-favoring residue (E), substitutions at position B2.5 have no effect on predicted fold. When b2b3.12 is fixed to a dimer-permissive residue (M), identity at position B2.5 strongly modulates the structural outcome (chemokine fractions 0–97%), demonstrating that b2b3.12 controls whether position B2.5 can influence conformational accessibility.

**Figure 3.**
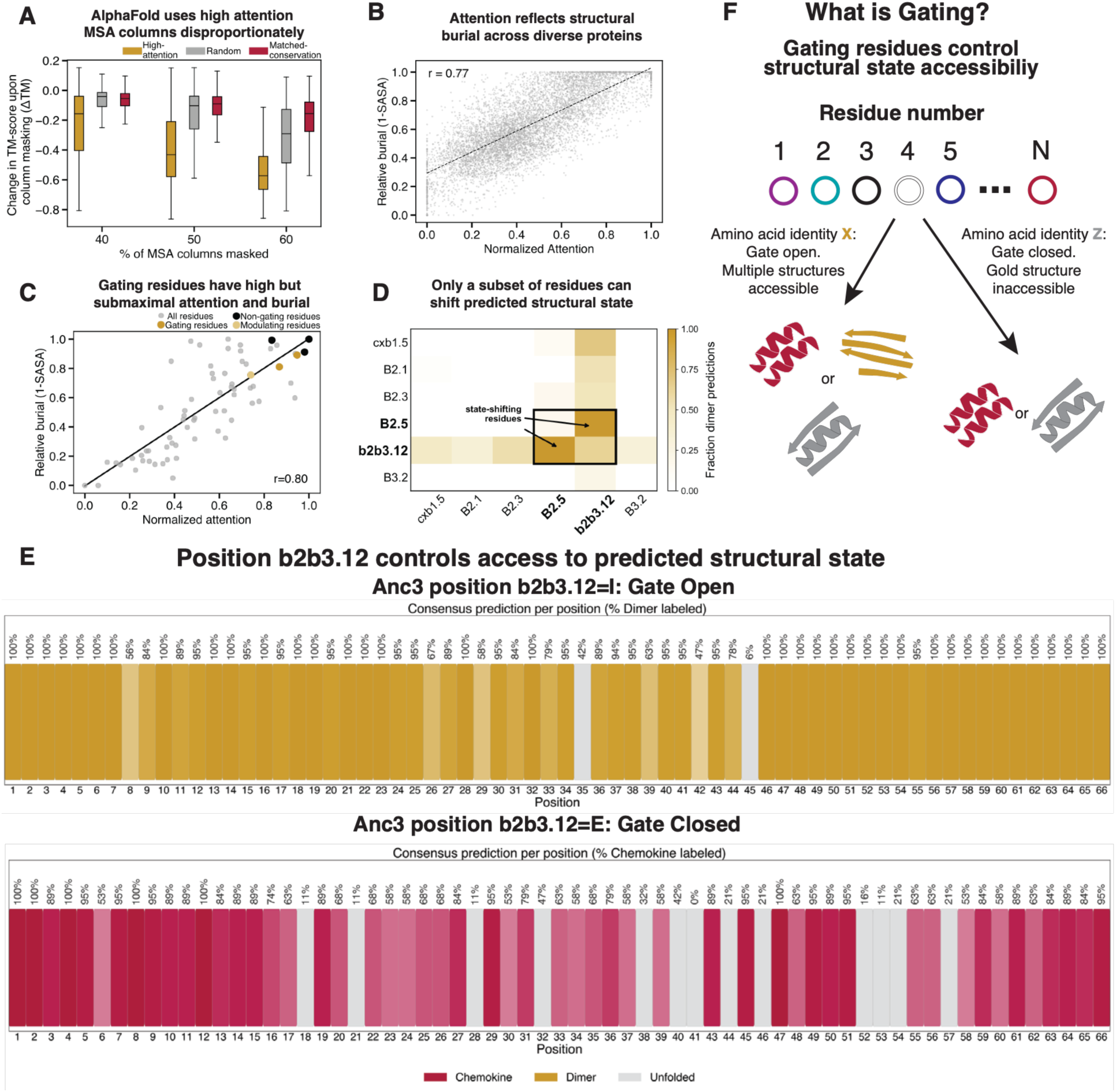
Attention reflects structural burial, but conformational accessibility is controlled by a sparse subset of gating residues. **(A)** Across 988 protein domains, high-attention MSA columns are disproportionately used during structural inference. Masking MSA columns corresponding to high-attention residues degrades AlphaFold predictions substantially more than random or conservation-matched masking across masking fractions of 40–60%, indicating that these positions are actively used during inference. **(B)** Across many sequence variants of 148 single-fold proteins, AlphaFold attention correlates strongly with residue burial (r = 0.77), indicating that the model preferentially focuses on buried positions that anchor protein structure. **(C)** Gating residues in XCL1 (gold) exhibit high but submaximal attention and burial relative to non-gating high-attention residues (black; positions B2.1, B2.3, and B3.2), demonstrating that attention and burial enrich for, but do not uniquely identify, conformational control points (r = 0.80). **(D)** Only a sparse subset of residues can alter the predicted structural state. Systematic pairwise mutagenesis of the six highest-attention positions shows that only positions B2.5 and b2b3.12 substantially shift the predicted conformation, whereas mutations at other highly attended positions have little effect despite comparable attention and burial. **(E)** Position b2b3.12 hierarchically controls access to predicted structural states. When b2b3.12 is fixed to a dimer-permissive residue (I), nearly all single mutants remain compatible with the dimer fold. When b2b3.12 is fixed to a chemokine-favoring residue (E), nearly all single mutants adopt the chemokine fold. Thus, the identity of a single residue determines which conformational states are predicted to be accessible across diverse mutational backgrounds. **(F)** Conceptual model of gating. A gating residue (residue 4 in diagram) is a conformational control point whose identity determines which structural states remain accessible. When the gate is open, multiple structural states remain accessible across many amino acid substitutions at sites other than the gate. Changing the identity of the gating residue suppresses access to the gold state, thereby restricting the range of conformations that AlphaFold predicts.

**Figure 4.**
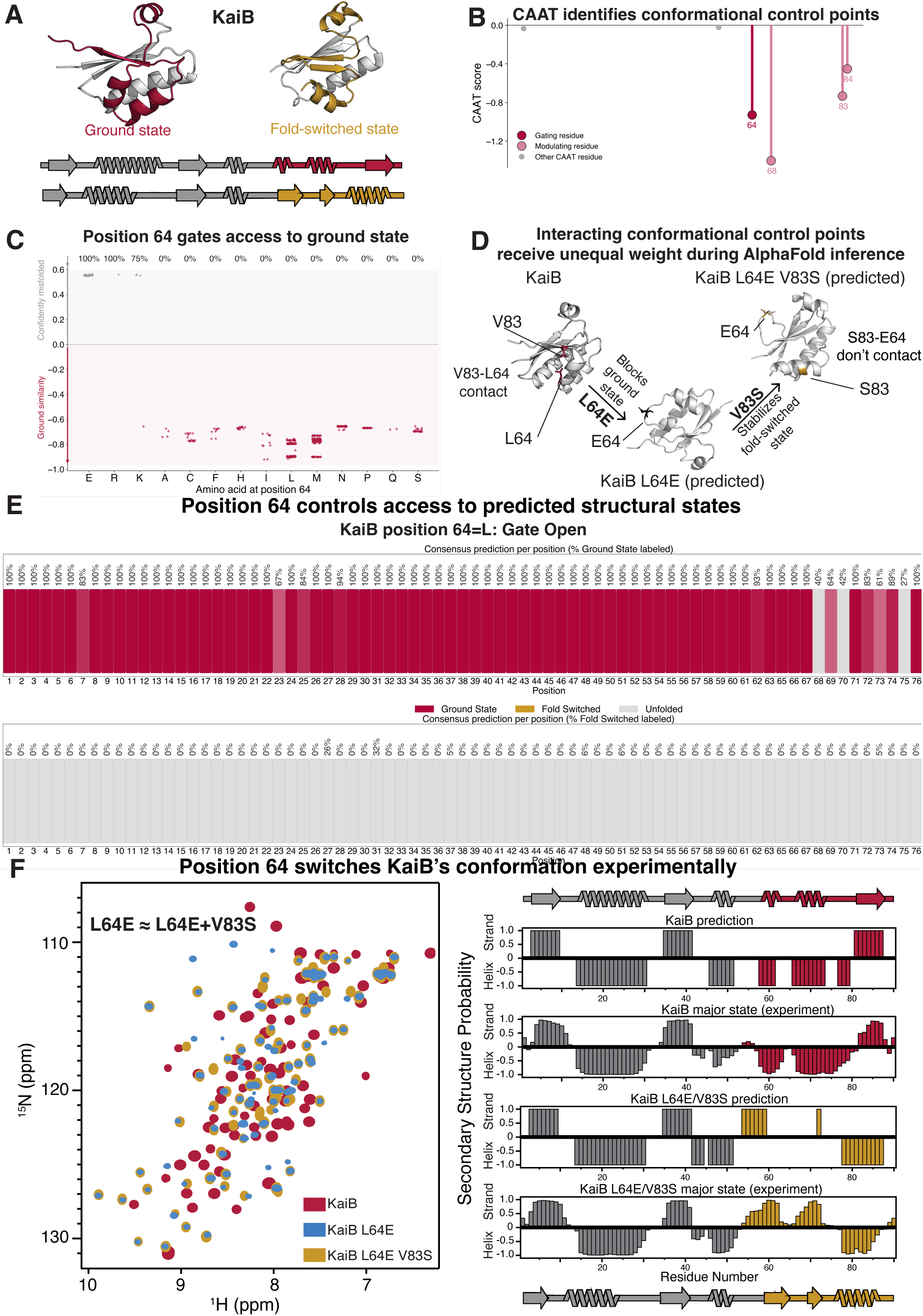
AlphaFold identifies biologically meaningful conformational control points but assigns them unequal weight during conformational selection. **(A)** KaiB adopts distinct ground-state (red) and fold-switched (gold; PDB 8FWJ, chain M) conformations that differ substantially in secondary-structure organization and regulate cyanobacterial circadian timing. **(B)** CAAT identifies a small set of conformational control points associated with structural outcome. Although L64 does not exhibit the strongest CAAT signal, it is AlphaFold’s dominant accessibility determinant. **(C)** Systematic mutagenesis of position 64 reveals conformational gating. Charged substitutions (E, R, K) abolish ground-state predictions, whereas most other amino acids retain access to the ground-state conformation. Ground-state similarity is shown relative to wild type. **(D)** Interacting conformational control points receive unequal weight during AlphaFold inference. In the fold-switched structure, L64 and V83 form a direct interaction. Substitution L64E disrupts this interaction and eliminates ground-state accessibility, whereas V83 primarily influences the identity of the accessible fold-switched state. These results indicate that AlphaFold disproportionately concentrates conformational control onto L64 despite the participation of both residues in the same interaction network. **(E)** Position 64 hierarchically controls access to predicted structural states. When L64 is retained (gate open), most single mutations throughout KaiB remain compatible with the ground-state conformation. By contrast, L64E (gate closed) globally suppresses ground-state accessibility across diverse mutational backgrounds, shifting predictions toward the fold-switched state. **(F)** Experimental validation by NMR spectroscopy. Substitution L64E shifts KaiB toward the fold-switched conformation, consistent with AlphaFold predictions. The experimentally observed secondary-structure profile of KaiB L64E/V83S closely matches the predicted fold-switched structure. Although V83S is sufficient to promote fold switching experimentally (**Fig S11B**), AlphaFold2 responds substantially more strongly to perturbation of L64, indicating that conformational accessibility is concentrated onto a dominant accessibility determinant despite experimental evidence that conformational control is distributed across interacting residues.

**Figure 5.**
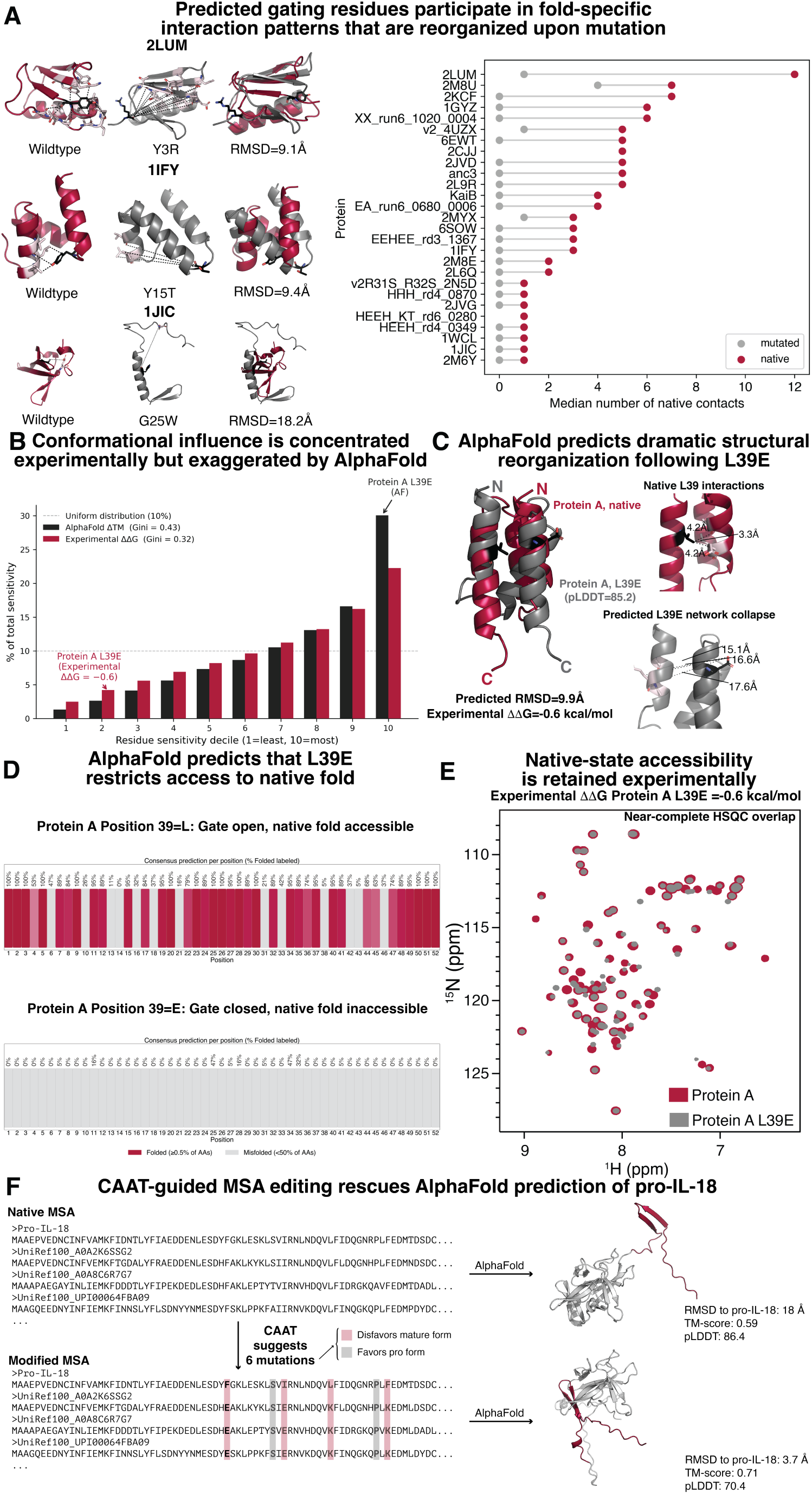
AlphaFold overweights conformational control relative to experiment. **(A)** Predicted gating residues participate in fold-specific interaction patterns that are reorganized upon mutation. Representative examples from diverse folds illustrate the structural consequences of gate-disrupting mutations identified by AlphaFold. Although the resulting structural rearrangements vary substantially among proteins, mutations frequently alter the interaction pattern associated with the predicted gating residue. For each protein, the median number of native contacts associated with the gating residue was compared with the number of those native contacts retained after mutation. In most cases, gate-disrupting mutations eliminated most or all native interactions associated with the predicted gating residue, although some mutations established new interaction patterns rather than simply removing contacts (**Figure S13**). These observations suggest that AlphaFold preferentially assigns conformational influence to residues whose interaction patterns differ strongly between predicted structural states. **(B)** AlphaFold assigns disproportionate conformational influence to a small subset of residues relative to experimental measurements. Residues were ranked by structural sensitivity (ΔTM, AlphaFold predictions) and experimental sensitivity (ΔΔG). AlphaFold concentrates approximately 30% of total predicted sensitivity into the highest decile, whereas experimental sensitivity is distributed substantially more broadly (Gini = 0.43 versus 0.32). Position L39 from Protein A, highlighted in red, falls within the highest sensitivity decile despite exhibiting a near-neutral experimental stability effect (ΔΔG = −0.6 kcal/mol). **(C)** AlphaFold predicts dramatic structural reorganization following mutation of a top-decile gating residue. Substitution L39E produces a high-confidence structural rearrangement (RMSD = 9.9 Å; pLDDT = 85.2) in which the helical packing geometry is reorganized. AlphaFold predicts that mutation disrupts the native L39 interaction network and eliminates contacts that stabilize the native structural arrangement. Native PDB structure: 6SOW. **(D)** AlphaFold predicts that L39 controls access to the native fold. When position 39 is fixed as leucine (native), most substitutions throughout the protein remain compatible with the native folded state. By contrast, when position 39 is fixed as glutamate, AlphaFold predicts that access to the native fold is globally suppressed across mutational backgrounds, indicating hierarchical gating behavior. **(E)** Experimental measurements do not support the predicted gate closure. Two-dimensional ^1^H–^15^N peak position diagrams of wild-type Protein A and the L39E variant exhibit near-complete spectral overlap despite the dramatic structural rearrangement predicted by AlphaFold, indicating retention of the native major conformational state. Thus, AlphaFold predicts loss of native-state accessibility where experimental data indicate that the native fold remains the major state. **(F)** Mechanistic interpretation enables targeted improvement of AlphaFold conformational prediction. CAAT was applied to both the pro- and mature forms of human interleukin-18 (IL-18) and identified six informative MSA columns: four whose residue identities favored the mature conformation (pink) and two whose identities could be replaced with alternatives predicted to favor the pro-form (gray). Editing these six columns substantially improved the predicted pro-form structure, reducing the RMSD to the experimental pro-form from 18 Å to 3.7 Å and increasing the TM-score from 0.59 to 0.71. Although the corrected prediction carried lower confidence (pLDDT = 70.4 versus 86.4), it more closely matched the experimentally observed structure. These results demonstrate that mechanistic interpretation of AlphaFold can identify MSA-level determinants of incorrect conformational predictions and that targeted modification of these determinants can substantially improve conformational prediction. The RMSD and pLDDT values differ slightly from those in Fig. 1A because AF3 was used for structure predictions here, whereas AF2 was used in 1A.

Together, these analyses show that AlphaFold represents conformational accessibility through a principle we term gating, in which surprisingly few residues hierarchically determine whether mutations elsewhere in the sequence influence structural outcome. Comparison with experiment reveals that AlphaFold captures this biologically meaningful architecture of conformational control while exaggerating its sparsity, explaining both its successes and its failures in predicting alternative conformations and mutational effects.

## Results

### AlphaFold2 internally encodes thermodynamic information but does not propagate it to structure predictions

We began by investigating which sequence features AlphaFold2 (AF2) utilizes to determine accessible conformational states, and to what extent those features correspond to true control points that govern protein folding. We chose to look at AF2’s attention weights because AF2 tends to predict alternative conformations more reliably than AlphaFold3 (AF3) (*9*). We utilized a dataset that reports protein stability measurements through cDNA display proteolysis on more than 500,000 mutations (up to 19 amino acid substitutions at many positions) across 148 proteins spanning diverse folds (*25*). This dataset enabled us to systematically compare experimentally measured mutational effects on protein stability with AF2’s internal representations and predicted structural outputs (**Fig. 1B**). These proteins were selected for the quality of their stability measurements and because AF2 accurately predicts their structures from single sequences, enabling analysis of mutational effects independent of MSA contributions. We first assessed AF2’s internal attention weights, which quantify the relative importance of residues during structure prediction, and binned residues by attention decile. To focus on features known to influence thermodynamic stability we compared the experimentally measured mutational effects on stability (ΔΔG) across the attention deciles. We observed a strong monotonic enrichment of destabilizing mutations (ΔΔG ≤ −3 kcal/mol) at high deciles (**Fig. 1C**). We also calculated the extent of burial (solvent accessible surface area, or SASA) in the predicted structure for each protein variant and plotted the values for each attention decile (**Fig. 1D**). Residue burial correlated with stability across these diverse proteins, explaining how AF2’s sensitivity to mutational effects could arise from training on experimental structures without an explicit energy function. The residues in higher attention deciles had the highest SASA values and the least variance. Together these results indicate that AF2 attention correlates with biophysical parameters that influence protein folding.

To determine how AF2’s internal representations propagate to its structure predictions, we measured the structural similarity between each mutant AlphaFold model and its cognate wild-type model using the template modeling score (TM-score (*26*)), and we defined ΔTM as the absolute change in TM-score, such that larger ΔTM values indicate larger predicted structural rearrangements. Across the attention deciles we found no systematic enrichment (r = −0.12, n = 10,322, **Fig. 1E**). High-attention residues were not more likely to produce structural changes when mutated. Since destabilizing mutations would typically be expected to decrease the equilibrium population of the native structure but not necessarily alter the structure itself, we examined the effects of mutations of prediction confidence as well. AF2’s confidence (pLDDT) also shows little relationship with high attention deciles (**Fig. S2**). Together, these results reveal a disconnect: AF2’s internal representations focus on features correlated with thermodynamic stability, yet this information is not obviously propagated to its predicted structures or confidence scores.

### AlphaFold represents conformational accessibility in XCL1 through a sparse hierarchy of gating residues

To further explore what sequence features AlphaFold uses to determine conformational access, we examined fold-switching proteins in depth. We began by investigating how mutations affect structure predictions using the human immunoprotein XCL1, which adopts two distinct conformations: a monomeric chemokine fold containing both an α-helix and β-sheets, and a dimeric β-sheet fold (**Fig. 2A**). These conformations perform distinct functions, with the monomer mediating GPCR signaling and the dimer binding fungal pathogens (*27, 28*). XCL1 is particularly well suited to probe conformational selection, as a panel of ancestral reconstructions and sequence variants has been experimentally characterized by ZZ-exchange experiments, revealing both single-folding and fold-switching variants (*22*). Running AlphaFold in single-sequence mode (predictions run with MSAs produce chemokine conformations exclusively), we predicted structures for all 19 experimentally characterized variants and compared them to experimental ground truth. Both AF2 and AF3 substantially overpredicted the dimer conformation. Across 25 replicates for each of 19 experimentally characterized variants, AF2 predicted the dimer conformation for 10 variants and AF3 for 8 variants, however only 4 of 19 were observed in the dimer state experiments (*22*) (**Fig. S3**). This discrepancy suggested that sequence variation in XCL1 could be used to interrogate features influencing AlphaFold predictions.

To identify features that drive AlphaFold’s predictive outcomes, we systematically examined the effect of sequence variation on predicted conformations. We use the common chemokine residue numbering system to account for alignment shifts among the 19 sequence variants (*29*). Although substitutions at most positions did not alter predictions, amino acid identity at positions cxb1.5, B2.5, and b2b3.12 strongly influenced the predicted fold. Chemokine predictions were associated with Y at position cxb1.5, T at position B2.5, and R at position b2b3.12, whereas dimer predictions were enriched for branched aliphatic residues (I, L, V) at these positions (**Fig. S3**). To test whether these positions alone determine AlphaFold’s conformational selection, we generated variants spanning combinations of amino acids at these sites. Both AF2 and AF3 predictions were largely determined by these identities: combinations containing branched aliphatic residues favored the dimer conformation, whereas polar and charged residues favored the chemokine fold (**Figs. 2B, S4**).

These results indicate that AlphaFold selects between XCL1 conformations using a small number of sequence determinants. Notably, introducing similar substitutions into CCL8, a single-folding homolog (35% sequence identity), induced analogous predicted conformational changes, suggesting that AlphaFold can apply this accessibility principle beyond fold-switching clades (**Fig. S5**). Together, these findings show that AlphaFold reduces high-dimensional sequence information to a small number of decisive features governing conformational accessibility of XCL1 and its ancestors. Because our earlier assessment revealed that AlphaFold’s internal representations correlate with parameters relevant for stability, we wondered if residues driving conformational selection would exhibit distinct representations corresponding to alternative folds.

To test this, we analyzed attention patterns used to predict folding of XCL1 and its variants as a proxy for AlphaFold’s internal information flow. We found interaction networks specific to the dimer conformation that were absent in sequences predicted to adopt the chemokine fold (**Fig. S6**). These differences were evident even between closely related sequences differing at only a few positions, indicating that attention patterns reflect sequence features associated with fold selection (**Fig. S7**). To generalize this analysis strategy, we developed an end-to-end pipeline that automates the generation of AlphaFold2 structures and analyzes the raw attention heads, comparing attention patterns across sequences. This framework, termed the Conformational Attention Analysis Tool (CAAT, **Fig. S8**), identified the three key positions governing XCL1 fold selection (**Fig. 2C**). Importantly, these positions correspond to an experimentally determined interaction network specific to the dimer fold (**Fig. 2A**). Notably, while our original discovery of the three critical residues required hundreds of AlphaFold predictions, CAAT requires only two AlphaFold runs followed by a few simple tensor operations, making it broadly applicable.

Among the three residues with highest CAAT scores, position b2b3.12 affected predictions strongly. To explore the possibility that changes at a single residue could influence conformational accessibility, we compared the similarity of structure predictions with every possible substitution at b2b3.12 to the chemokine and dimer folds using TM-score. Amino acid identity at this position strongly modulated the predicted conformation (**Fig. 2D**), with the fraction of dimer predictions ranging from 0% (D, E, K, Q, S) to 88% (I). The biophysical properties of amino acids enabled us to infer the basis for this pattern: hydrophobic residues favored the dimer conformation, while polar and charged residues favored the chemokine fold, consistent with the distinct packing environments of position b2b3.12 in the two structures (**Fig. 2A**). This basis is consistent with position b2b3.12 being buried in the dimer conformation’s interface. These results suggest that the identity of the amino acid at this position limits conformational accessibility, consistent with XCL1 experiments showing that b2b3.12 is hydrophobic in all dimer-forming variants.

To determine the relationship between individual residues that govern conformational prediction, we examined the ability of substitutions at B2.5, which also has a high CAAT score, to affect conformational prediction in the presence of a chemokine favoring or dimer-permitting amino acid at position b2b3.12 (**Fig. 2E**). When position b2b3.12 was fixed to glutamate — a chemokine- favoring residue — substitutions at position B2.5 had no effect: and all variants were predicted to adopt the chemokine fold. By contrast, when position b2b3.12 was fixed to methionine — a dimer- permissive residue — the identity of position B2.5 strongly modulated the structural outcome, with dimer fractions spanning 3–100%. Together, these results demonstrate that XCL1 fold selection is governed by a hierarchical gating process in which the identity of position b2b3.12 determines whether other residues can influence the predicted fold.

### AlphaFold attention identifies structurally important residues in XCL1, but only a sparse subset governs conformational accessibility

Having established that a small number of residues govern fold selection in XCL1, we investigated structural and biochemical properties that might distinguish these gating positions from other residues. MSAs are also a potential source of conformational control. To test whether high-attention residues are causally used by AlphaFold during structural inference — rather than merely correlated with important positions — we measured the ΔTM of predictions upon MSA column knockout on 988 protein domains with different folds (**Methods**). Systematically masking MSA columns corresponding to high-attention residues degraded AlphaFold predictions substantially more than masking randomly selected columns or conservation-matched controls (**Fig. 3A**). This effect was consistent across masking fractions of 40%, 50%, and 60%, indicating that high-attention positions are not passive correlates of structural importance but are actively used by the model during inference.

We wondered whether the associations between attention and residue burial (**Fig. 1D**) might also be relevant for gating because buried residues have greater potential for key stabilizing interactions. We utilized the same dataset with >500,000 mutations across 148 single-fold proteins with different topologies. Here, instead of breaking out attention deciles, we plotted the relative burial against normalized attention for each protein in the dataset. Attention remained strongly correlated with residue burial (r = 0.77; **Fig. 3B**), indicating that AlphaFold’s preferential focus on buried positions is a general architectural property rather than a consequence of the stability filter applied in Figure 1.

To test whether burial alone determines conformational accessibility we looked again at XCL1. We calculated and plotted the SASA of each residue relative to normalized AlphaFold attention (**Fig. 3C**, r=0.80). The gating residues (cxb1.5, B2.5, b2b3.12; gold) had high but submaximal attention and burial, while several other residues (positions B2.1, B2.3, B3.2; black) had higher attention and/or burial than the gating positions.

To directly test which positions can shift the predicted structural state, we performed systematic pairwise mutagenesis of six high-attention positions in XCL1 (cxb1.5, B2.1, B2.3, B2.5, b2b3.12, B3.2) and measured the fraction of dimer predictions across amino acid combinations. Only mutations at positions B2.5 and b2b3.12 shifted the predicted structural state from chemokine to dimer; mutations at the other positions — despite having comparable or higher attention and burial — had negligible effects on predicted conformation (**Fig. 3D**). The specificity of state-shifting to\ positions B2.5 and b2b3.12 confirms that gating is a property of a sparse subset of residues, not a general feature of buried or highly attended positions. Importantly, previous experiments indicate that both B2.5 and b2b3.12 are required for XCL1 variants to assume the dimer conformation (*22*).

To assess the scope of b2b3.12’s predicted conformational control, we ran AlphaFold2 predictions on all possible single mutants at every other position in XCL1 and identified the predicted fold at each site. Strikingly, when b2b3.12=I, nearly every variant was predicted to assume the dimer fold (**Fig. 3E**). By contrast, when b2b3.12=E, nearly every variant was predicted to assume the chemokine fold (**Fig. 3E**). Thus, the identity of a single amino acid determines which conformations are accessible at inference. Importantly, AlphaFold3 predicts that b2b3.12 exhibits the same gating behavior, indicating that sparse control of XCL1’s conformational accessibility spans different AI architectures (**Figure S9**). Gating therefore refers to hierarchical control of conformational accessibility, in which the identity of residues at one or more positions determines whether the effects of mutations elsewhere in the sequence can influence structural outcome.

Together, these results refine the mechanistic model of how AlphaFold selects between conformations. AlphaFold concentrates attention on buried residues across diverse proteins, and this attention is causally used during structural inference. Within the set of high-attention residues, however, only a sparse subset—gating residues—controls access to structural states. As illustrated in the conceptual model (**Fig. 3F**), the sequence background permits access to multiple potential conformations, whereas the identity of a gating residue determines which of those structural states remain accessible. Changing the identity of a gating residue can therefore suppress access to one or more conformational states while other states remain accessible.

### AlphaFold identifies biologically meaningful determinants of conformational accessibility but assigns them unequal influence

To determine whether gating represents a general organizational principle of conformational control, we examined the fold-switching protein KaiB from *Rhodobacter sphaeroides*. KaiB adopts distinct ground-state and fold-switched conformations that differ in secondary structure organization and regulate the periodicity of the cyanobacterial circadian clock (**Fig. 4A**) (*23, 30*). Applying CAAT to KaiB identified a small set of residues associated with conformational outcome, including L64 and V83 (**Fig. 4B**). Through extensive sampling, a previous study found that mutating three of these amino acids simultaneously—I68R, V83D, N84A—switched KaiB’s conformation from ground to fold-switched (*24*), but, to our knowledge, L64 had not previously been implicated or experimentally tested. Systematic *in silico* mutagenesis revealed that position 64 gates KaiB’s ground state. Whereas wild-type KaiB is predicted to adopt the ground-state fold, substitution of L64 by charged amino acids (E, R, K) eliminated ground-state predictions and shifted AlphaFold toward an alternative structural state (**Fig. 4C, D**).

Among the residues identified by CAAT, L64 exerted hierarchical rather than additive control over predicted conformational accessibility. Retention of L64 preserved ground-state accessibility across most mutational backgrounds, whereas L64E almost completely abolished ground-state accessibility irrespective of mutations elsewhere in the protein (**Fig. 4E**); this result holds for AF3 as well, though it predicts the fold-switched conformation for L64E rather than a conformation that is partially ground state and partially fold-switched (**Figure S10**). Thus, AlphaFold organizes conformational control hierarchically, with the identity of a single residue determining whether an entire structural state remains accessible to subsequent sequence variation.

L64 and V83 are physically interacting residues within the ground state structure, yet perturbation analysis revealed distinct mechanistic roles for these positions within the AlphaFold model. L64 hierarchically gates access to the ground state — its retention preserves ground-state accessibility across most mutational backgrounds, whereas L64E almost completely abolishes it regardless of mutations elsewhere in the protein (**Fig. 4E**). By contrast, V83 primarily influences which fold-switched state is accessible rather than whether the ground state can be reached (**Figs. 4D, E**). This asymmetry predicts that perturbation of L64 should have substantially larger effects on conformational outcome than perturbation of V83 despite their direct interaction — a prediction AlphaFold makes strongly, with far greater response to L64E than to V83S (**Fig. S11A**).

With XCL1, experimental studies and CAAT analysis identified the same residues capable of determining conformational accessibility experimentally. We hypothesized that the gating principle used by AlphaFold may be a bona fide biophysical principle governing protein folding. To explore this possibility, we investigated the effects of mutations on KaiB folding using nuclear magnetic resonance (NMR) spectroscopy.

NMR results revealed that both L64E and V83S individually shifted KaiB toward the fold-switched state, as did the double mutant L64E/V83S (**Fig. 4F, S11B**), consistent with AlphaFold’s identification of both positions as conformational determinants. The experimentally observed secondary-structure profile of KaiB L64E/V83S agreed closely with the predicted fold-switched conformation. However, V83S alone was sufficient to promote fold switching experimentally (**Fig. S11A**), whereas AlphaFold assigned substantially greater influence to L64E than to V83S (**Fig. S11B**), indicating that conformational control is distributed across both interacting residues experimentally but concentrated disproportionately onto L64 within AlphaFold’s representation. Together, these results indicate that AlphaFold identifies biologically meaningful determinants of conformational accessibility but concentrates their influence on some gating residues more strongly than observed experimentally. Whether compensatory mutations can restore access to the gated conformation remains an open question, as our experimental validation focused on gate-closing rather than gate-reopening mutations.

We also examined a third fold-switching protein, the C-terminal domain (CTD) of RfaH (*31*), to further investigate the relationship between CAAT identified gating residues and experimentally measured conformation accessibility. CAAT identified L142 as the strongest determinant of the high-confidence RfaH CTD helical-state prediction which is consistent with an experimentally characterized RfaH CTD minor state (*32*) (**Fig. S12A**) The substitution L142S caused AlphaFold predictions to collapse from the single helical structure to a heterogeneous ensemble of predominantly low-confidence conformations, indicating loss of a well-defined folded state (**Fig. S12B**). Circular dichroism measurements confirmed that RfaH L142S is unfolded (**Fig. S12C**), indicating that CAAT correctly identified an experimentally meaningful destabilizing residue. In addition, these experiments revealed that CAAT-identified conformational control points can regulate access not only to alternative folds but also to the folded state itself, which may be relevant for single-fold proteins.

### AlphaFold overweights conformational control of single-fold proteins relative to experiment

To investigate whether hierarchical gating residues also govern conformational accessibility in single-fold proteins, we used AF2 to predict the structures of mutants from the dataset of >500,000 experimentally measured mutational effects across 148 proteins (*25*). Focusing on a subset of unique folds, we identified gating residues as positions where single mutations consistently altered predicted structure relative to wildtype. By this approach, 28 gating residues were identified across diverse folds, nearly all of which participate in fold-specific interaction patterns that are frequently altered by mutation (**Fig. 5A**). In most cases, mutations at gating residues eliminate native interactions associated with the gating position, although some mutations establish alternative interaction networks (**Fig. S13**). These predicted gating residues were significantly enriched for experimentally destabilizing mutations relative to all measured mutations (median ΔΔG = −2.0 versus −0.3 kcal/mol; KS test p = 5 × 10⁻¹⁴; **Fig. S14**), indicating that mutations to putative gating residues have genuine energetic importance. Nevertheless, energetic importance and conformational control are not always proportional: AlphaFold preferentially weights structurally distinctive positions within fold-specific interaction architectures, yet structural distinctiveness does not guarantee dominant energetic control.

To assess whether AlphaFold accurately weights individual conformational control points relative to their experimental importance, we returned to the dataset of >500,000 experimentally measured mutational effects across 148 proteins (*25*). Residues were ranked by AlphaFold-predicted structural sensitivity (ΔTM) and grouped into deciles; experimental stability measurements (ΔΔG) were analyzed in parallel (**Fig. 5B**). Although both prediction and experiment concentrated sensitivity in a subset of residues, AlphaFold’s distribution was more concentrated, with the highest sensitivity decile accounting for ∼30% of total predicted structural change. Importantly, predicted and experimentally sensitive residues did not always agree: projecting experimental data onto the AlphaFold-defined ranking revealed that experimental sensitivity was distributed substantially more broadly, with the top decile accounting for less than half of the influence AlphaFold assigned to it for both ΔTM and ΔpLDDT (**Fig. S15**). These results indicate that AlphaFold concentrates conformational influence into fewer residues than experimental measurements suggest, and that the residues AlphaFold predicts as most influential do not necessarily align with experimental observation.

Position 39 in Protein A serves as an example. AlphaFold identifies L39 as one of the strongest conformational control points in the protein in the highest sensitivity decile (**Fig. 5B**). In fact, substitution L39E is predicted to induce a dramatic structural reorganization despite a near-neutral experimentally measured stability effect (ΔΔG = −0.6 kcal/mol). AlphaFold2 predicts inversion of the protein’s helical packing arrangement, yielding a high-confidence structure that differs substantially from the native state (RMSD = 9.9 Å; pLDDT = 85.2; **Fig. 5C**); AlphaFold3 predicts the same inverted structure in response to L39E. Structural inspection suggests that this prediction may arise because L39 participates in a native interaction network that is lost in the rearranged structure.

Consistent with this interpretation, AlphaFold identifies position 39 as a gating residue. When position 39 is fixed as leucine, most mutational backgrounds remain compatible with the native fold. By contrast, when fixed as glutamate, position 39 globally suppresses native-state predictions across the sequence, indicating that AlphaFold predicts that L39E closes access to the native conformational state (**Fig. 5D**).

We tested this prediction experimentally using NMR spectroscopy. Contrary to AlphaFold’s prediction, a two-dimensional ^1^H–^15^N peak position diagram of wild-type Protein A and the L39E variant exhibited near-complete overlap, indicating retention of the native major conformational state (**Fig. 5E**). This observation is consistent with the previously measured near-neutral stability effect of L39E (ΔΔG = −0.6 kcal/mol). Thus, AlphaFold predicts loss of native-state accessibility in a case where experimental measurements indicate that the native fold remains both accessible and nearly unchanged energetically.

The Protein A example demonstrates that AlphaFold can substantially overestimate the conformational influence of individual residues even when they occupy structurally privileged positions and are associated with experimentally important interactions. This distinction provides a mechanistic explanation for why AlphaFold can both identify biologically meaningful conformational control points and misrepresent their influence.

AlphaFold can also miss genuine gating residues by assigning them insufficient attention, producing the opposite failure mode. In a fold-switched stabilized variant of *Thermosynechococcus elongatus* KaiB, four of the five mutations required to switch the fold occupy low-attention positions (**Fig. S16A**), and AlphaFold predicts both the wildtype and mutant forms assume the ground state, failing to detect the experimentally observed fold switch (*23*). Similarly, AlphaFold predicts no conformational effect for XCL1 R23A/R43A, whereas ¹⁹F NMR experiments show that this double mutation shifts the dominant state from dimer to monomer (**Fig. S16B**). Both mutations occupy low-attention positions. Finally, AlphaFold predicts that a low-attention mutation to RfaH (H152L) preserves the minor helical state, whereas NMR experiments show no evidence for this state and differential scanning calorimetry indicates substantial stabilization of the β-sheet conformation (**Fig. S16C**). Together these examples demonstrate that AlphaFold systematically underweights low-attention positions, missing experimentally important conformational determinants even when their effects are large.

### Mechanistic interpretation of AlphaFold enables targeted improvement of pro-IL-18 conformational prediction

Having established why AlphaFold can weight some conformational control points improperly, we next asked whether this mechanistic understanding could improve incorrect conformational predictions. Our earlier finding that AlphaFold uses high-attention MSA columns disproportionately during structural inference (**Fig. 3B**) suggested that CAAT could be extended to identify specific MSA columns whose residue identities bias predictions toward incorrect conformational states. We applied this approach to human pro-interleukin-18 (pro-IL-18), which AlphaFold confidently predicts in the mature conformation despite the pro-form adopting a distinct experimental structure (AF3: RMSD = 18 Å; TM-score = 0.59; pLDDT = 88.4, **Fig. 5F**; AF2 metrics similar to AF3 in **Fig. 1A**).

We leveraged CAAT to recover predictions similar to pro-IL-18 without prior knowledge (**Fig. 5F**). We projected AlphaFold’s attention weights onto a simplified input MSA combining the pro-IL-18 sequence with a single sequence generated by ProteinMPNN (*33*). This approach repurposes ProteinMPNN as a probe of AlphaFold’s conformational logic: by generating a sequence predicted to adopt the incorrect fold, it implicitly identifies residues that anchor the mispredicted conformation, providing a principled basis for selecting MSA edits without prior knowledge of the target structure. CAAT identified four columns whose residue identities favored the mature conformation and two columns whose residue identities favored the pro-form conformation (**Figure S17**). Editing these six MSA columns substantially improved AF3’s pro-form prediction, reducing the RMSD from 18 Å to 3.7 Å (4.1 Å for AF2) and increasing the TM-score from 0.59 to 0.71 for both AF3 and AF2 (**Fig. 5F**). The highest-confidence prediction from the modified MSAs carried lower confidence than the original incorrect prediction (pLDDT = 70.4 versus 86.4), consistent with the interpretation that AlphaFold’s high confidence in the original incorrect prediction reflected strong emphasis on MSA features associated with the mature conformation. All 15 AF2 predictions generated from this modified MSA assumed conformations similar to the pro-form, demonstrating that CAAT-guided edits produced consistent predictions across AlphaFold models and seeds. These results demonstrate that CAAT can identify MSA-level determinants of incorrect conformational predictions and that targeted modification of these determinants can substantially improve prediction accuracy, extending the conformational control framework from single-sequence predictions to MSA-dependent conformational inference.

## Discussion

Our results suggest that conformational accessibility is often organized around surprisingly few gating residues that hierarchically control access to alternative structural states. Mechanistic interpretation of AlphaFold revealed this organizational principle while also exposing how the model systematically exaggerates it. To identify gating residues, we developed the Conformational Attention Analysis Tool (CAAT), which extracts conformational control points from AlphaFold’s internal representations.

A defining feature of gating is its hierarchical dominance: AlphaFold predicts that once a gating residue closes access to a conformation, no single mutation elsewhere in the sequence can restore it. This prediction has been validated in the gate-closing direction — mutations at predicted gating residues shift experimentally observed conformational equilibria as predicted. Whether compensatory mutations could reopen a closed gate in physical proteins remains to be tested and represents a key prediction of the gating framework. Supporting this prediction, all experimentally characterized XCL1 variants that assume the dimer fold had branched aliphatic residues in predicted gating positions, consistent with gate-open states, whereas the conformations of all variants with polar or charged residues at these positions were observed to assume to the chemokine fold only, consistent with closed gates (*22*). Together, these findings indicate that AlphaFold captures a biologically meaningful architecture of conformational control that frequently corresponds to experimentally important determinants of protein behavior.

At the same time, AlphaFold frequently underweights other experimentally important determinants. AlphaFold failed to predict conformational switching in a fold-switched stabilized variant of *T. elongatus* KaiB (*23*), a double mutant of XCL1 that shifts the dominant state from dimer to monomer, and a mutant of RfaH that stabilizes the β-sheet conformation (**Fig. S16**). In each case, experimentally important mutations received comparatively little emphasis in AlphaFold’s internal representation. AlphaFold also did not identify the V83S mutation in KaiB from *R. sphaeroides* as a gate, though the mutation was sufficient to flip its fold (**Fig. S11**). Conversely, AlphaFold occasionally assigned dominant influence to residues whose experimental effects were modest, as illustrated by Protein A L39E. Together, these observations suggest that AlphaFold represents conformational control through a sparse hierarchy of residues that control conformational accessibility. When experimentally important residues are strongly emphasized within this hierarchy, AlphaFold can successfully predict conformational effects. However, residues that receive little emphasis may exert substantial experimental influence without altering AlphaFold predictions, while overemphasized residues can produce confident but experimentally unsupported conformational predictions.

The examples examined here–combined with the statistical analysis of ΔΔG values for 30 predicted gates–suggest that sparse conformational control points are a genuine feature of many proteins. However, AlphaFold does not appear to weight these control points according to their energetic contribution to conformational equilibria (*34, 35*). Instead, AlphaFold preferentially assigns influence to residues occupying structurally distinctive positions within fold-specific interaction architectures. Such positions often correspond to experimentally meaningful determinants of conformational accessibility, but geometric distinctiveness does not necessarily imply dominant energetic control. Physical factors such as electrostatics, solvation, temperature, pH, and conformational entropy contribute to conformational equilibria but are not explicitly represented in AlphaFold (*36*). Consequently, AlphaFold can miss experimentally important mutations whose effects arise primarily through these mechanisms, such as the electrostatically driven R23A/R43A switch in XCL1 (**Fig. S16**).

These findings have implications beyond structure prediction. If sparse conformational control points are a general organizational feature of protein structure, they may represent privileged targets for engineering conformational switches (*37, 38*), designing allosteric regulators (*39*), and understanding how evolution diversifies protein function (*40*) by reprogramming access to alternative conformational states. The identification of gating residues through mechanistic interpretation of AlphaFold provides a tractable strategy for discovering these determinants, while the distortions we have characterized suggest opportunities to build models that represent conformational accessibility more faithfully. For instance, systematizing approaches like the one used to recover the experimentally characterized conformation of pro-IL-18 could lead to more robust predictors of alternative conformations and mutational effects (*41*), an area where AlphaFold and related methods currently struggle (*8, 17, 42*). Our approach combines two AI tools — ProteinMPNN and AlphaFold via CAAT — in a manner neither was designed for, using one model’s output to interrogate another model’s internal representations and correct its predictions, suggesting a general strategy for recovering conformations that AlphaFold confidently mispredicts. More broadly, this work illustrates that mechanistic interpretation can move beyond explaining AI models (*43–47*) to discovering biological principles and improving their predictive capabilities. Similar approaches may prove valuable for interpreting and improving AI models applied to other problems of interest.

## Methods Summary

AlphaFold2 predictions were performed using ColabFold (*48*) version 1.5.5 with all five models; AlphaFold3 predictions used version 3.0.0. Attention weights were extracted from AlphaFold2’s Evoformer block during inference using jax.experimental.io_callback. We used AlphaFold2’s weights rather than AlphaFold3’s because AlphaFold2 tends to predict alternative conformations more robustly than AlphaFold3 (*9*). CAAT scores are computed from AlphaFold’s attention weights by comparing attention patterns between sequences predicted to adopt alternative conformations, weighted by amino acid similarity (BLOSUM62). Gating residues in fold-switching proteins were identified by systematic *in silico* mutagenesis and confirmed by NMR spectroscopy. For single-fold proteins, gating residues were identified across 28 representative folds from a dataset of >500,000 experimentally measured mutational effects across 148 proteins (*25*); two fold-switching proteins (XCL1 and KaiB) were included in the list of 30. Conformational sensitivity was calculated by averaging the three most structurally perturbative mutations per position and normalizing within each protein. MSA column knockout experiments were performed on 988 CATH domains to assess causal use of high-attention positions during inference. CAAT-guided MSA editing was applied to human pro-interleukin-18 by identifying and modifying MSA columns whose residue identities were inconsistent with the pro-form conformation. NMR experiments were performed on Bruker spectrometers at 600 or 800 MHz; backbone assignments used standard triple-resonance experiments. Circular dichroism measurements were made on a Chirascan Q100. Full computational and experimental methods are provided in the supplementary materials.

## Acknowledgments

We thank Marius Clore, Attila Szabo, Keir Neuman, and Eugene Koonin for helpful discussions and Dan Herschlag and George Rose for critically reading the manuscript. This research was supported in part by the Division of Intramural Research at the National Library of Medicine, National Institutes of Health (NIH, LM202011, L.L.P.). The contributions of the NIH authors are considered Works of the United States Government. The findings and conclusions presented in this paper are those of the authors and do not necessarily reflect the views of the NIH or the U.S. Department of Health and Human Services. This work utilized the computational resources of the NIH HPC Biowulf cluster (https://hpc.nih.gov). Certain software is identified in this paper in order to specify the experimental procedure adequately. Such identification is not intended to imply recommendation or endorsement of any product or service by NIST, nor is it intended to imply that the materials or equipment identified are necessarily the best available for the purpose.

## Data and Code Availability

Code and data can be found at: https://github.com/prameshsharma25/CAAT

