## Supplementary Information for "AlphaFold reveals but sometimes distorts an organizational principle of protein folding"

**The PDF file includes:**

Materials and Methods
Figs. S1 to S17
References 48-66

### Materials and Methods

AlphaFold predictions

AlphaFold predictions were performed with ColabFold (*48*) version 1.5.5 for AlphaFold2 with alphafold2 weights and AlphaFold3 (*11, 12*) version 3.0.0. Predictions from both architectures were performed using all 5 models.

Attention heads

Attention head weights exist only as intermediate tensors within AlphaFold2's (AF2's) JIT-compiled JAX computation graph and are not exposed as model outputs, so they cannot be retrieved by simply calling the model and reading a return value. To access them without modifying AF2's source code, we used jax.experimental.io_callback, which allows a Python callback with side effects to be executed during trace execution, giving us a reliable way to read out intermediate values as the model runs without altering the underlying computation. This required deactivating AF2's custom memory optimization, which otherwise breaks each attention head into smaller pieces distributed across the GPU for enhanced parallelization; with this optimization active, attention heads could not be reassembled and localized within the model, precluding downstream interpretability analysis. Attention heads were extracted from each run of ColabFold (all 5 models, 3 recycles) by deactivating AF's custom memory optimization, allowing entire attention heads to be saved chronologically from the GPU, via the jax.experimental.io_callback module, as the model gets executed. This approach causes a slowdown in the runtime of AF, but we believe this is the optimal tradeoff between simplicity and efficiency that does not significantly impact the utility of CAAT. There are two primary sources of slowdown. The first is due to turning off AF's memory optimization, which breaks each attention head into smaller pieces for enhanced parallelization. We remove this optimization because it obfuscates the location of each attention head in the model, inhibiting the interpretability of our results. The second source of slowdown comes from the time cost of transferring attention head values from the GPU to the CPU during execution. For different computational experiments with greater memory requirements, more sophisticated memory management may be required.

Attention analysis

Attention tensors with shape nx4xnxn (n=number of residues in the target protein; 4= number of attention heads) were selected at each network layer, and softmax renormalization was applied along the final dimension, retroactively, to match the softmax within AF2. Axes 0, 1, and 2 were summed, producing a one-dimensional tensor (vector) of length n. Min-max normalization was performed on this vector, and the resulting scaled average was used for downstream analysis. Anc0 and XCL1 sequences were aligned in biopython (*49*) and BLOSUM62 (*50*) scores were obtained for all aligned amino acids. BLOSUM62 score was multiplied by the average attention previously obtained to determine position-specific CAAT scores. Negative scores (those specific to the conformation of interest) were plotted to obtain the amino acids important to XCL1 and Anc0 in **Figure 2C.** Attention values were systematically lower at N- and C- termini than in the center of the protein. To avoid false positives in CAAT in cases where aligned sequences had gaps at termini, attention differences between the first/last 5 amino acids after/before a terminal gap were ignored.

Prediction of effect of mutations on stability

All proteins from Tsuboyama2023_Dataset2_Dataset3_20230416 that had mutations with calculated ddG values in the "ddG_ML" column were used for the attention analysis. Structures of these wildtype sequences were initially predicted with ColabFold single-sequence (5 seeds, 3 recycles, 5 models). The predictions were compared against their corresponding experimentally determined structures using TM-align (*26*). Proteins retained for our analysis met the following criteria: 80% of their single-sequence predictions had TM-scores ≥ 0.75 and pLDDTs ≥ 70. This left 148 unmutated proteins with 566,415 mutations. CAAT was run on each unmutated sequence, and ColabFold (5 models, 1 seed) was run on all mutations. TM-align was used to compare all mutated predictions to the unmutated reference structures to calculate ΔTM scores.

To determine if AlphaFold gave higher attention to positions that caused destabilizing mutations, all mutations that resulted in experimentally measured ΔΔG <= -3 were counted and assigned their respective attention values for the unmutated position. To determine the impact of the mutations on ColabFold predictions, the AlphaFold2 model that resulted in the greatest change in ΔΔG was selected (top 1 out of 5 models) for each mutation, and the change in TM-score from unmutated, and the change in pLDDT were plotted against the respective attention value for the position that was mutated.

SASA calculations

SASA calculations were made on AF2 rank 1 predictions using FreeSASA (*51*). Values were normalized by relative accessible surface area values from (https://www.science.org/doi/10.1126/science.4023714). Calculated relative SASA values greater than 1 were min-max normalized.

XCL1 predictions

ColabFold and AF3 were initially run on the sequences of XCL1 and its homologs in single sequence mode, 3 recycles. Unlike AF2, AF3 was trained after Anc0 was added to the training set (PDB 7JH1, released 12/30/2020). Anc0 differs from other variants by a deletion and a glutamine mutation at position 28, which likely altered some of AF3's predictions. Anc2j had the deletion but not the mutation. An N28E mutation shifted all its predictions to dimer.

For the strip plots in Figure S3, all predictions were scored using TM-align. Reference structures had PDB IDs 1J9O for chemokine and 2JP1 for dimer. For 2JP1 residues 1-50 were used for TM-align, ignoring predictions of a spurious C-terminal helix noted previously (*52*). Strip plot points were colored with the requirement that TM-scores relative to experiment exceeded 0.5. If TM-scores against both reference structures were < 0.5, then the point was colored black. Only points with plDDT > 72 were plotted. The same approach was used for the AF3 strip plots in S1, but with all points were plotted regardless of plDDT.

For the scatter plots in **Figure 2B**, predictions were run on all 19 variants with a Cb1b2.12G mutation added, following the experiments from Dishman, et al. (*22*). Anc0's likely addition to AF3's training set made AF3 more sensitive Anc0-specfic sequence modifications. Accordingly, the sequences of Anc2c, Anc2j, and Anc2l were also modified to contain the N28E ΔA29 mutations for AF3 runs only. This restored the expected patterns in response to the three mutations cxb1.5, B2.5, and b2b3.12: chemokine fold for charged/polar; dimer fold for branched aliphatic. Minimal exceptions were found in these patterns. For both AF2 and AF3 all combinations of I,L,V resulted in dimer predictions for all variants except for I/L/V cxb1.5, I/V B2.5, L b2b3.12, which produced the chemokine fold in Anc0 only (6/17965 models). For AF2, charged residues in positions cxb1.5, B2.5, and b2b3.12 caused some unphysical dimer predictions in variants with sequence identities closer to XCL1 when K, T, H, N were in position cxb1.5 or N was in position B2.5. Removing these cases, 25 dimer predictions remained out of 17,965. All had S or T in positions B2.5 or b2b3.12. These predictions, which represent 0.2% of predictions are not shown in Figure 2B. Predictions were generated by making all I,L,V combinations in positions cxb1.5, B2.5, and b2b3.12, all 5 AF2 models, 3 random seeds. Charged residue predictions were made by randomly sampling 1000 combinations of amino acids in positions cxb1.5, B2.5, and b2b3.12 except for the patterns mentioned above, all 5 AF2 models, 3 random seeds. For AF3, the same chemokine prediction for Anc0 was observed for I/L/V cxb1.5, I/V B2.5, Lb2b3.12 as in AF2; these points, representing < 1% of all dimer predictions, were not shown in Figure S4. Otherwise, no exceptional sequence rules were observed for AF3 predictions. Figure S4 shows all AF3 predictions with plDDT scores ≥ 62 and **Figure 2B** shows AF2 predictions with plDDT scores ≥ 70. The same trend holds for AF3 predictions with plDDT scores ≥ 70, but few dimer predictions were made with predictions above this threshold.

XCL1 gating

For **Figures 2D and 2E**, ColabFold 1.5.5 was run with 3 seeds, 3 recycles, and all 5 models for all 19 XCL1 variants, with all sequence positions held fixed at wildtype identities except where noted. For **Figure 2D**, position b2b3.12 was sampled across all 20 amino acids, generating 5,700 structures total. For **Figure 2E**, position b2b3.12 was fixed at M (gate open) or E (gate closed) while position B2.5 was sampled across all 20 amino acids, generating 5,700 structures per panel. For **Figure 3D**, all 20 amino acids were sampled at each of the six highest-attention XCL1 positions, and the three substitutions most perturbative to the native fold (largest change in ΔTM relative to chemokine in a 20 amino acid scan across each protein) at each position were identified and exhaustively sampled as pairwise combinations. The top three substitutions were: cxb1.5: I, L, V; B2.1: A, G, I; B2.3: I, L, V; B2.5: G, I, V; b2b3.12: I, L, V; B3.2: I, L, M. Because this analysis was intended to assess access to the dimer fold, only variants not predicted to access the dimer conformation from wildtype sequences were included: Anc0, Anc2, Anc2a, Anc2b, Anc2c, Anc2d, Anc2f, Anc2i, and Anc2l.

For **Figure 3E**, all 19 possible substitutions were made at every residue position of XCL1 Anc3, except position b2b3.12 (residue 42), which was held fixed at I or E for the two panels respectively. ColabFold was run with 3 seeds, 3 recycles, and all 5 models, generating 21,375 models total. Models with pLDDT ≥ 70 were classified as folded; a minimum TM-score of 0.55 was required for classification as either the chemokine or dimer fold, with the slightly lower threshold reflecting XCL1's disordered terminal regions. We calculated the %dimer/chemokine for every mutation took the consensus percentage for each position.

Number of contacts

Biopython's NeighborSearch was used to calculate the number of contacts on AF rank 1 predictions. All heavy atoms from backbone and side chain were used in calculating the number of contacts. Residues within 5 Angstroms were considered in-contact and all residues further away were excluded. Residues within 4 amino acids on either side were also excluded. For number of contacts to the gating residue, 8 amino acids on either side were also excluded.

Role of MSAs and attention

All representative topology domains from the CATH database (*53*) were downloaded from the latest release v4_3_0. The following filters were applied to remove domains: less than 50 residues, greater than 300 residues, 80% single sequence predictions with TM-scores less than 0.75, 80% MSA predictions with TM-scores less than 0.75, greater than 50% disordered based on DSSP. The filtering steps resulted in 988 CATH domains remaining. For the MSA knockouts, columns that received the highest attention were mutated to X, gaps were maintained, and the target sequence remained unmutated. Conservation scores were calculated based on Henikoff calculations. For the high attention columns, the corresponding conservation scores were binned, 0-0.3, 0.3-0.7, and 0.7-1. Columns that were removed due to greater than 50% gaps based on Henikoff were assigned -1. The distribution of scores within each bin was recorded, and low attention columns were selected to obtain the same distribution of conservation scores. If the same distribution could not be obtained with the corresponding low attention columns, the lowest possible attention columns possible were added until the distribution was obtained.

XCL1 attention

CAAT was run on single sequences of XCL1 to obtain the average attention values for each position. Free SASA was used to calculate SASA. Each amino acid's SASA value was normalized by the standard ASA values. Two amino acids (T and V) had normalized SASA values above 1, so all T and V were min-max normalized. Number of contacts was calculated with BioPython's NeighborSearch using all heavy atoms.

KaiB gating

For **Figure 4B**, gating calculations were run with Colabfold1.5.5, 50 seeds, 12 recycles all 5 models for the KaiB *R. Sphaeroides* sequence (MGRRLVLYVAGQTPKSLAAISNLRRICEENLPGQYEVEVIDLKQNPRLAKEHSIVAIPTL VRELPVPIRKIIGDLSDKEQVLVNLKMDME). We used 50 seeds to ensure statistical robustness: we sampled sequence variation in only 1 sequence for KaiB rather than the 19 XCL1 variants; likewise, 12 recycles were required to consistently generate accurate, high-confidence KaiB predictions. Position 64 sampled all 20 amino acids while the rest of the sequence positions stayed fixed at wildtype residue identities, totaling 4,017 predictions. For **Figure 4E**, all 19 mutations were made at all residue positions of KaiB, except for position 64, which was held fixed at L and E for the two panels, respectively. For each sequence variant, ColabFold was run with 3 seeds, 12 recycles on all 5 models for a total of 25,365 models. Only models with pLDDT scores ≥ 70 were considered folded, with a minimum TM-score of 0.6 required to be classified as either the ground or fold-switched state.

KaiB Rs and variants

[$^{15}$N]-enriched samples were expressed in the same fashion as described above. Uniformly [$^{2}$H/$^{13}$C/$^{15}$N]-enriched *R. sphaeroides* KaiB L64E;V83S was prepared as follows. Multiple colonies were picked and grown in 3 mL LB media (Research Products International) dissolved in 99.8% $D_2O$ (Cambridge Isotope Laboratories) and grown overnight at 37 °C. The next day, The 2.5 mL overnight was diluted into 22.5 mL modified M9+ medium(*54*) with 3 g/L [$^{2}$H,$^{13}$C] glucose and 1.5 g/L $^{15}NH_4Cl$ in 99.8% $D_2O$ and grown for 24 hours at 37 °C. The next day, the culture was diluted to 1 L modified M9+ and at $OD_{600}$= 0.8, the culture was induced with 0.3 mM IPTG in $D_2O$. After 7 hours of expression at 37 °C, the cells were harvested as described above.

Cells were resuspended, lysed, underwent nickel affinity purification, SUMO cleavage, and secondary nickel affinity purification as described above, except TCEP was added to nickel affinity buffers. After, the sample was diluted 5-fold in 20 mM MES, 0.5 mM TCEP, pH = 5.5, and then purified via cation-affinity chromatography using a home-made SP-resin column (Cytiva). The protein was isolated over a 5 CV linear gradient from 0-40% 20 mM MES, 2 M NaCl, 0.5 mM TCEP, pH = 5.5. Pure fractions were determined by SDS-page gel analysis, pooled, and purity was validated by mass spectrometry. Protein concentrations assessed using absorbance at 205 nm (*55*).

NMR Spectroscopy

NMR measurements were acquired on Bruker Avance III 600 MHz, Avance Neo 600 MHz, or Avance III 800 MHz spectrometers equipped with z-gradient cryogenic probes. Spectra were processed using NMRpipe (*56*). KaiB measurements were made in 25 mM MOPS, 50 mM NaCl, 2 mM TCEP, 1 mM PMSF, 5% $D_2O$, pH = 6.5 at 35 °C. Backbone assignments of KaiB L64E;V83S were performed at 600 uM concentration, 35 °C using TROSY-HSQC, HNCO, HNCA, HNCACB, HN(CO)CA, HN(CO)CACB experiments. Non-uniform sampled data was reconstructed using SMILE (*57*) and all data processed using NMRPipe. Assignments performed using CCPN (*58*). TALOS+ was used for secondary structure prediction (*59*). RfaH CTD and CTD H152L measurements made in 20 mM potassium phosphate, 100 mM NaCl, 1 mM ethylenediaminetetraacetic acid, 10% (v/v) $D_2O$, pH 6.5 at 23 °C.

RfaH constructs

The RfaH C-terminal Domain (*31*) was ordered from Twist Biosciences (San Francisco, CA) and cloned into a pPal7 vector with an N-terminal 6x his-tag and bdSUMO solubility tag (*60*). H152L and L142S variants were generated using site-directed mutagenesis and confirmed by whole-plasmid sequencing. ecNusG CTD, KaiB RS and all described variants were ordered from Genscript (Piscataway, NJ) and cloned into pET28a vectors with an N-terminal 14x his-tag and bdSUMO solubility tag. All plasmids were transformed into *E. coli* BL21 (DE3) cells (Agilent).

RfaH purification

[$^{15}$N]-enriched protein expression was based on previous methods (*61, 62*). 2.8L flasks containing 1 L of Terrific Broth (Research Products International) were autoclaved. From a Luria Broth(LB)/Agar plate + 100 mg/mL ampicillin or frozen cell stock, a 25 mL Terrific Broth + 100 mg/mL ampicillin or 50 mg/mL kanamycin overnight culture was inoculated and grown at 37 °C. The next day, 1 L Terrific Broth flasks + 100 mg/mL ampicillin or 50 mg/mL kanamycin were inoculated with 2.5 mL overnight culture. At $OD_{600}$ = 0.6, flasks were spun at 6,000 rcf, 4 °C, for 15 minutes. Supernatant was poured off and cells resuspended in half of the original volume of 1X M9 salts, pH 7.4. Cells were again pelleted at 6,000 rcf, 4 °C, for 15 minutes, and then resuspended in 1/4 the original volume, in sterile filtered 2X M9 salts, 4 g/L D-glucose, 1 g/L $^{15}NH_4Cl$, 1X Modified Eagles medium, 2 mM $MgSO_4$, 100 mg/mL ampicillin or 50 mg/mL kanamycin, 5 mL of 200 X trace element solution (*63*). Unenriched protein was prepared in a similar fashion, except the overnight culture was used to inoculate 0.5 or 1 L Terrific Broth media + 100 mg/mL ampicillin or 50 mg/mL kanamycin and shaken at 180 rpm at 37 ºC to an $OD_{600}$ = 0.6. Cells were transferred to a shaker at 18 °C, and shaken for 30 minutes, after which, Isopropyl β-D-1-thiogalactopyranoside (IPTG) was added to a final concentration of 200 mM.

The next day, cells were harvested by centrifuging at 6,000 rcf, 4 °C, for 15 minutes. Cells were resuspended in 50 mM tris(hydroxymethyl)aminomethane (tris), 150 mM NaCl, 20 mM imidazole, 1 mM dithiothreitol, 5% glycerol, pH 8.8 at 4 °C. ½ cOmplete protease inhibitor tablet (Roche) was added, along with 100 U Benzonase Nuclease (Millipore). Cells were lysed (Microfluidics, M110P), spun at 38,724 rcf, 4 °C for 45 minutes, and then syringe filtered (0.45 mm).

Filtered lysate was then loaded on HisTrap HP (Cytiva) column, washed with 50 mM $KP_i$, 100 mM NaCl, pH = 7.4, then eluted from the column with a 4 CV linear gradient to 20% elution buffer (50 mM $KP_i$, 100 mM NaCl, 500 mM imidazole, pH 7.4), followed by immediate elution 100% elution buffer. Fractions containing protein of interest were pooled and 100 ng bdSUMO protease was added, and the sample dialyzed overnight at 4° C in wash buffer with 0.2 mM tris(2-carboxyethyl)phosphine) (TCEP) added. The next day, the cleaved sample was loaded again on the HisTrap column and washed, collecting the flow-through. The flow-though was concentrated and then further purified using size exclusion chromatography, pre-equilibrated in 100 mM $KP_i$, pH 7.4 (HiLoad Superdex 75 pg, Cytiva). Pure fractions were determined by SDS-page gel analysis, pooled, and purity was validated by mass spectrometry (6230 ESI-TOF LC/MS, Agilent). Protein concentrations assessed using absorbance at 205 nm (*55*).

Circular dichroism measurements

Measurements were made on a Chirascan Q100 (Applied Photophysics) in 1 mm quartz cuvettes (Hellma) in 100 mM Potassium Phosphate, pH 7.4. Concentration of RfaH CTD L142S was 17.3 mM, and the concentrations of RfaH CTD, RfaH CTD H152L, ecNusG CTD were 20 mM. Data was collected in 1 nm step sizes, 1 nm/s scan rate, at 20 °C. 10 scans were made for each protein, with measurements averaged. Triplicated buffer blanks were averaged and subtracted. Spectra were then converted to Molar Residue Elipicity $[\theta]_{MRE}$ using equation 1:

$$[\theta]_{MRE} = \frac{\theta * \varepsilon}{L * N * A}$$

Where $\theta$ is the measured ellipticity, $\varepsilon$ is the molar absorbtivity at 205 nm estimated from https://spin.niddk.nih.gov/clore/Software/A205.html (*55*), L is the cuvette pathlength, N is the number of amino acids, and A is the measured absorbance at 205 nm (Nanodrop One, Thermo Scientific).

Gating residues

Among the 148 wildtype proteins that met our criteria from the stability dataset, 30 unique representative folds were identified using FoldSeek (*64*). Potential gates were identified in 28 cases by identifying all single point mutations that resulted in the largest changes in median TM-scores across all 5 models. The gate was tested by performing all single point mutations for all positions. Gates were determined a success if less than 25% of the predictions were folded with the gate mutated. The number of residues in contact with the gating residue was determined with KDball. The total number of contacts for native and mutated, as well as the change in contacts from native were measured.

Conformational sensitivity

For each single mutant across the 148 proteins, five AlphaFold2 predictions were performed using all five models with one seed and three recycles. The median change in TM-score (ΔTM) or pLDDT relative to wildtype was calculated across the five predictions and assigned to the corresponding residue position. To calculate predicted structural sensitivity for each position, the three mutations producing the largest median ΔTM (or ΔpLDDT) were averaged and normalized by the sum of all per-position averages within each protein. Three mutations were used to balance sensitivity to large effects while reducing the influence of outliers. Experimental sensitivity was calculated analogously: for each residue position, the three mutations with the largest absolute change in stability (|ΔΔG|) were averaged and normalized by the sum of all per-position averages within each protein.

Protein A gating

For **Figure 5E**, gating calculations were run with Colabfold1.5.5, 50 seeds, 3 recycles all 5 models for Protein A. All 19 mutations were made at all residue positions of Protein A, except for position 39, which was held fixed at L and E for the two panels. For each sequence variant, ColabFold was run with 3 seeds, 3 recycles on all 5 models for a total of 14,535 models for each figure panel. Only models with pLDDT scores ≥ 70 were considered folded, with a minimum TM-score of 0.6 required to be classified as native.

Recovering the pro-IL-18 fold

The incorrectly predicted structure of pro-IL-18 generated by ColabFold was passed into ProteinMPNN to generate 1 sequence using a sampling temperature of 0.1. This sequence–39% identical to pro-IL-18–was aligned below the pro-IL-18 sequence target as a 2-sequence MSA passed into ColabFold. A test run with 1 seed and 12 recycles confirmed that this MSA consistently produced structures similar to the incorrect prediction, with TM-scores ranging from 0.79-0.84. This two-sequence MSA was used instead of a full MSA to pinpoint sequence differences between the target and MPNN sequences that may engender different structure predictions. CAAT was applied to the two sequence MSA with 12 recycles and all other settings as described in the Attention Methods sections. High attention regions were focused on the first 72 residues, reasoning that the first 36 residues comprise the IL-18's pro-domain and extending the analysis window to twice the pro-domain length to allow for potential conformational influence beyond the insertion itself. This approach assumes that the MSA features governing the incorrect prediction are concentrated near the pro-domain insertion rather than distributed throughout the sequence, which was sufficient to recover a native-like structure. Using this approach, CAAT identified the top 6 highest attention positions. Substitutions at these positions were chosen based on conservation between the pro-IL-18 and ProteinMPNN sequences. Four residues were conserved between the two sequences and hydrophobic (F38, V47, L56, F66); we reasoned that these residues anchor the incorrect mature conformation and therefore mutated the corresponding MSA columns — but not the target sequence itself — to charged residues (F38E, V47E, L56K, F66K). Two residues were not conserved between pro-IL-18 and the ProteinMPNN sequence; we reasoned that these may encode information favoring the correct pro-form and therefore mutated the MSA columns to match the pro-IL-18 identity (S46, P64). This combination of six MSA edits produced 15/15 AF2 predictions and the highest-confidence AF3 prediction resembling the experimentally determined pro-IL-18 conformation. Structural similarities were calculated using TM-scores and RMSDs with 2 refinement cycles in PyMOL against the experimentally determined pro-IL-18 structure (PDB ID: 8URV). Subsets of 5 or fewer mutations to MSA columns produced less accurate and/or less consistent predictions.

XCL1 protein preparations

Human XCL1(1-72) protein, which lacks the disordered C-terminal tail (residues 73–93), was expressed in DL39(DE3)E. coli, using a chemically defined media supplemented with 100 mg/L of 4-fluorophenylalanine and all 19 other naturally occurring amino acids as described (*65*). The inclusion body pellet containing hXCL1(1-72) was refolded and purified as previously described (*66*). Incorporation of a single fluorine atom was verified by electrospray mass spectrometry and presumed to reflect the substitution of F39, the only phenylalanine in the human XCL1 sequence. $^{19}F$ NMR spectra of 4-F-Phe labeled hXCL1(1-72) (100 µM) in aqueous buffer containing 20 mM NaPO4, 50 mM NaCl, pH 6.5 were acquired on a Bruker Avance III 500 NMR spectrometer at 35 ˚C using a spectral width of 37500 Hz, 256 scans, and 8192 total data points. Trifluoroacetic acid (100 µM) was added to each sample and used as an internal standard for normalization of peak intensities. Chemical shifts referencing was to an external trichloro-fluoro-methane (0 $^{19}F$ ppm).

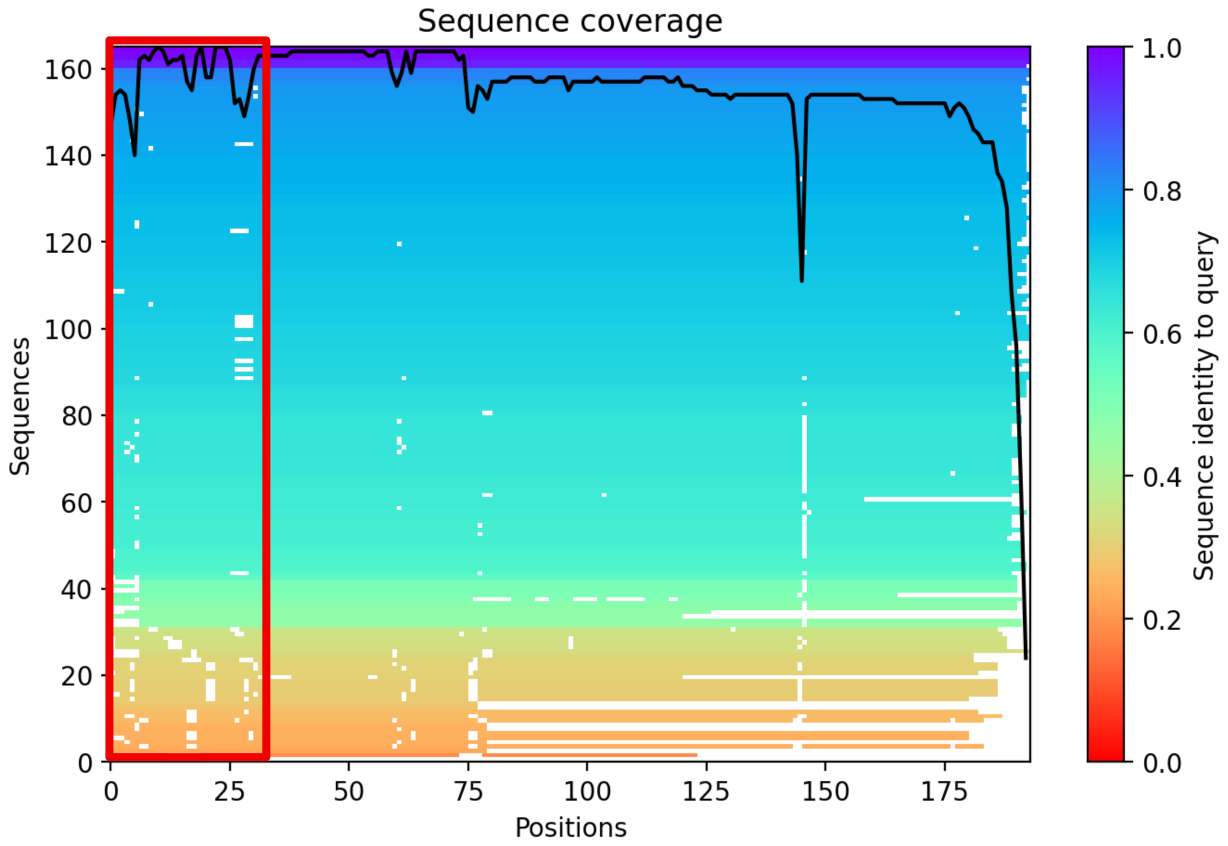


**Figure S1. AlphaFold confidently predicts the mature conformation of human interleukin-18 despite having adequate sequence coverage of the 36-residue NTD insertion from the pro-form.** Sequences with at least 80% gaps in the first 36 residues were removed from the pro-form MSA. The MSA still contains sequences with varying sequence identity.

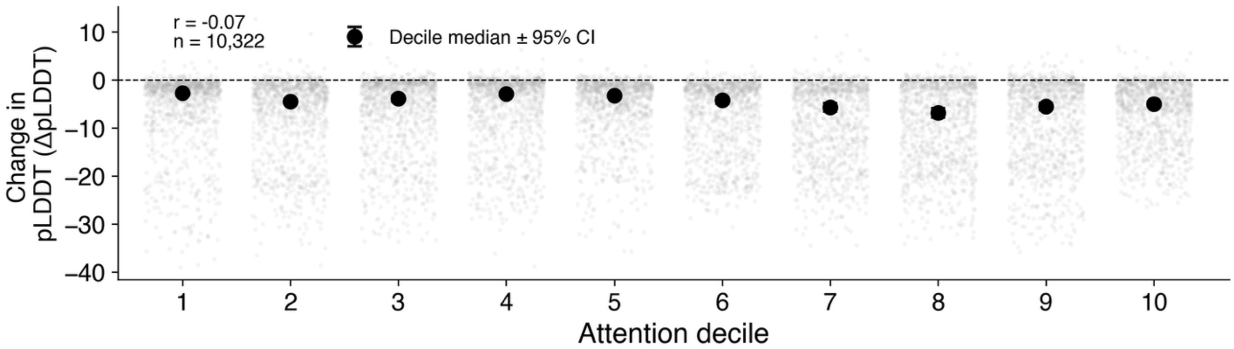


**Figure S2. AlphaFold's confidence does not reflect stability information.** AlphaFold's changes in confidence (ΔpLDDT) relative to wildtype show no relationship with attention decile (r = -0.07, n = 10,322), indicating that high-attention residues are not more likely to produce confidence changes when mutated. Further, pLDDT is minimally perturbed, though each mutation in the dataset destabilizes the protein by a minimum 3 kcal/mol.

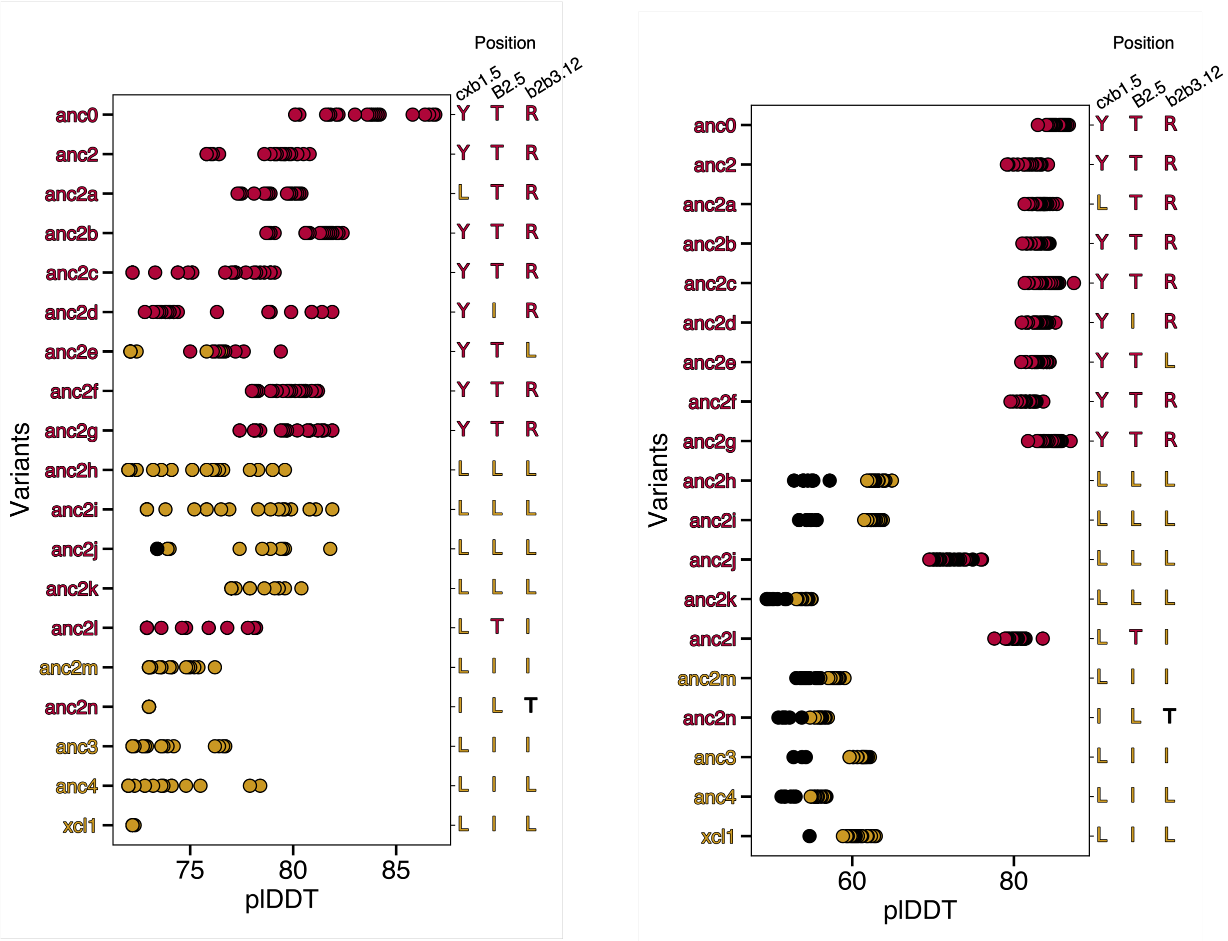


**Figure S3. AlphaFold overpredicts the dimer conformation (gold circles among XCL1 variants.** Red names were experimentally observed to assume the chemokine fold only; gold names exchanged between dimer and chemokine. Anc2e, 2h, 2i, 2j, and 2k are all predicted to assume the dimer conformation, which was not detected experimentally. Positions cxb1.5, B2.5, and b2b3.12 of these proteins' aligned sequences are enriched in Y, T, R, respectively for the chemokine conformation, and L, I for the dimer conformation.

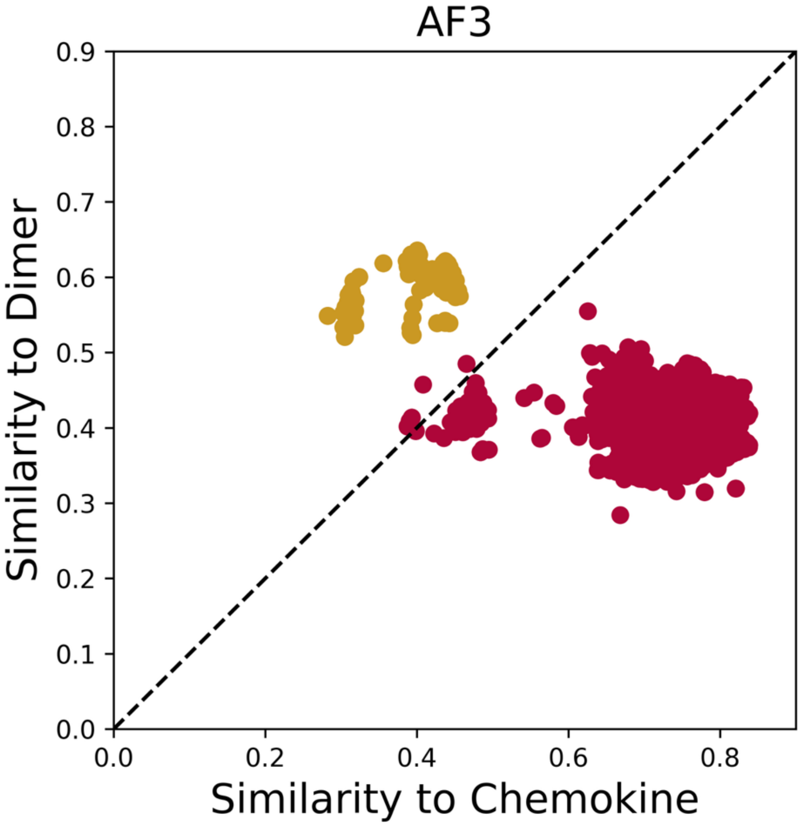


**Figure S4. Three XCL1 positions govern AF3's conformational selection of XCL1.** When the three CAAT amino acids are branched aliphatic (yellow) only the dimer conformation is predicted confidently; when the same three amino acids are charged/polar (red), the chemokine conformation is predicted confidently. Here, we use pLDDT of 62 as confidence threshold, but pLDDT = 70 yields a similar result but with fewer confident dimer predictions.

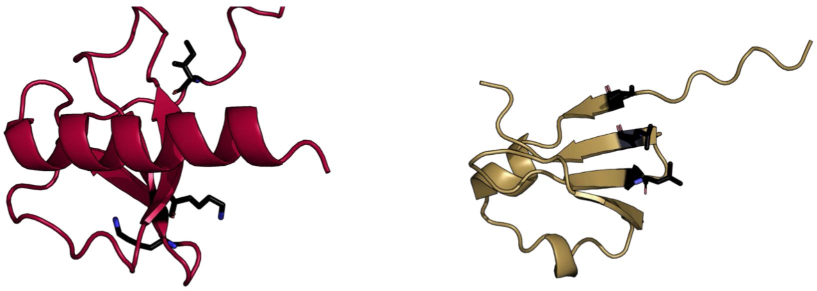

**CCL8 (Experiment)**

**CCL8 AlphaFold prediction cxb1.5 (I), B2.5 (L), b2b3.12 (L)**

**Figure S5. AF2 predicts that mutations to I/L in the same 3 positions cause CCL8 to assume the dimer fold.** This is unlikely since it is outside of the XCL1 clade. PDB ID of experimentally determined MCP-2: 1ESR. Nevertheless, the top-ranked AF2 prediction of MCP-2, had a pLDDT of 77.1 after 12 recycles. Like the other XCL1 variants, position Cb1b2.12 was mutated to G.

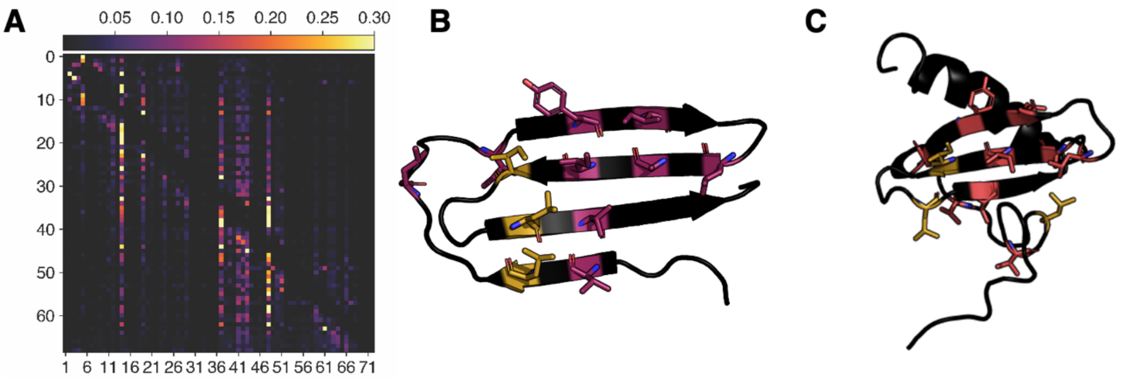


**Figure S6. AF2 pays attention to the dimer interaction network when predicting the dimer fold.** A. Attended amino acids correspond to vertical lines in the attention head; in this case layer 6, head 0, residue 1 was used. B. The attended amino acids in A are mapped onto XCL1's dimer structure and correspond to an interaction network; positions cxb1.5, B2.5, and b2b3.12 are highlighted in gold; the other attended positions are purple. C. The interaction network in the XCL1 attention head and in the dimer fold is not present in the chemokine fold; positions cxb1.5, B2.5, and b2b3.12 are highlighted in gold; the other attended positions are orange.

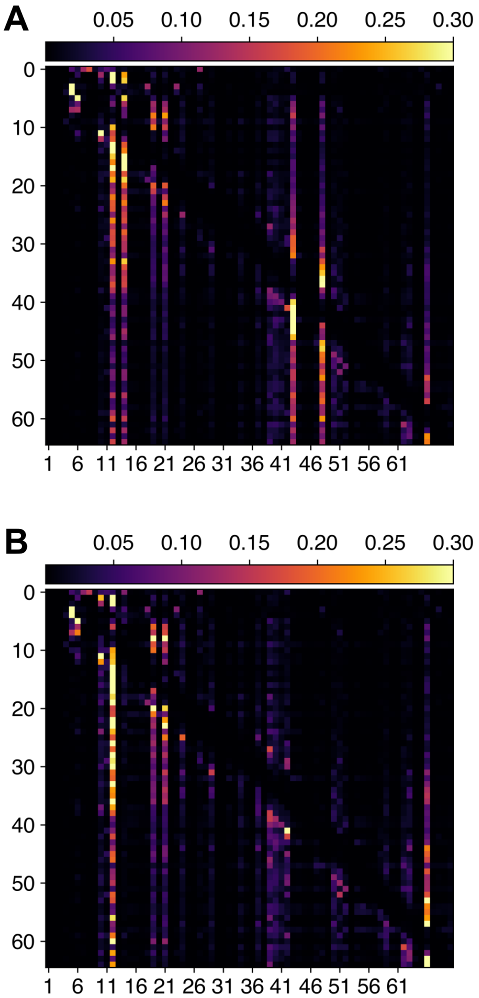


**Figure S7. AF2 attention patterns for Anc2f cxb.15, B2.5, and b2b3.12 and wildtype Anc2f resemble those of XCL1 and Anc0, respectively.** A. The attention pattern of Anc2f cxb.15, B2.5, b2b3.12 corresponds to an interaction network unique to the XCL1 dimer fold. Note that positions cxb1.5, B2.5, and b2b3.12 are lit up as they are for XCL1 in Figure 2B. AF2 predicts that Anc2f cxb.15, B2.5, b2b3.12 assumes the dimer fold. B. The attention pattern of wildtype Anc2f does not correspond to the interaction network unique to the XCL1 dimer fold. Note that positions cxb.15, B2.5, and b2b3.12 are not lit up as they are for XCL1 in Figure 2B; rather the attention pattern of Anc2f resembles Anc0. AF2 predicts that Anc2f assumes the chemokine fold. Attention patterns were taken at residue 0, head 0, layer 6, recycle 0.

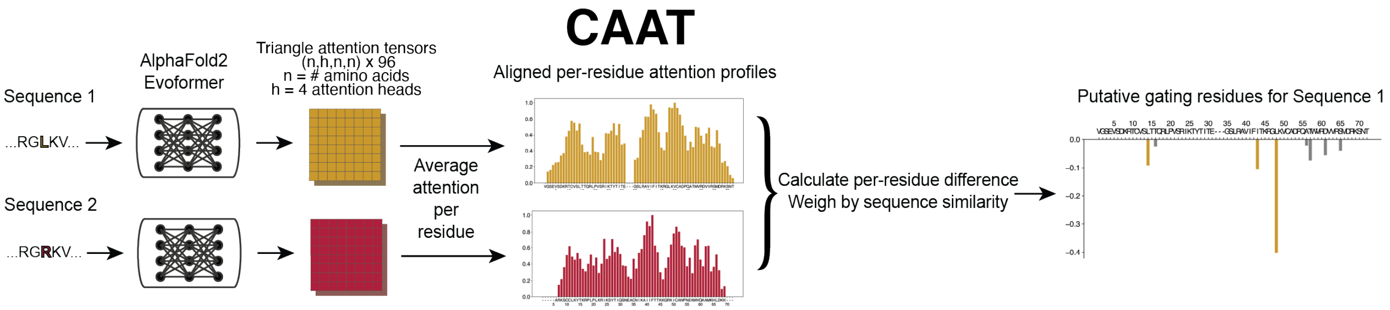


**Figure S8. A schematic of the CAAT algorithm.** Attention was averaged across all network layers for all recycles.

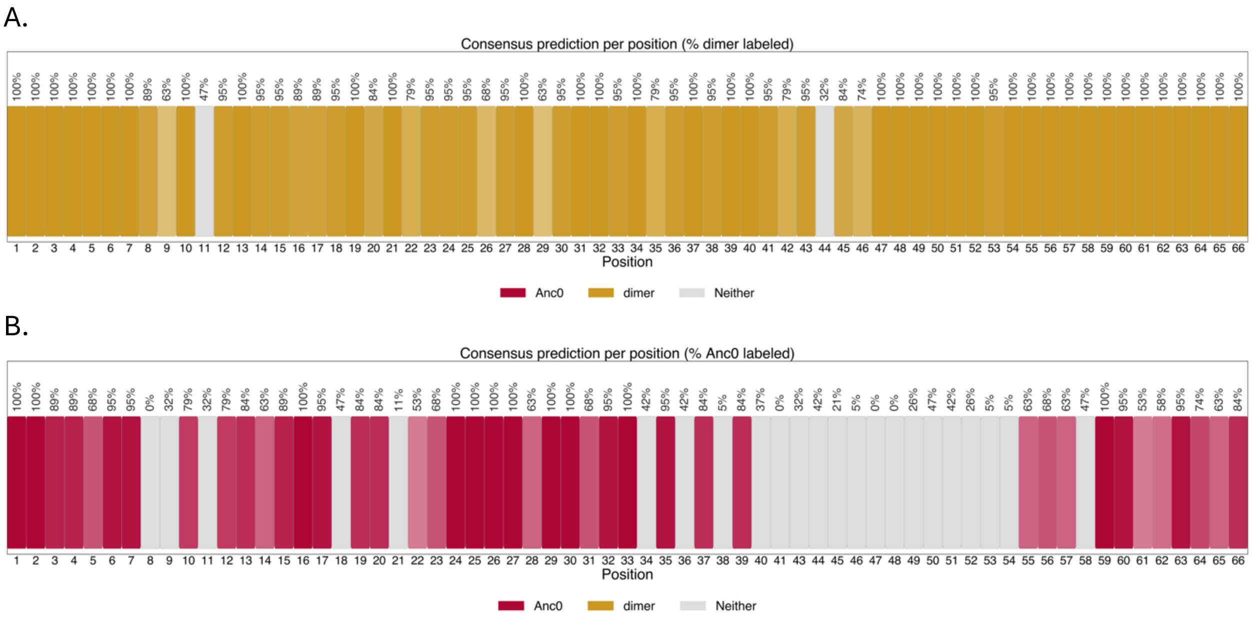


**Figure S9. AlphaFold3 exhibits the same gating behavior for XCL1 anc3 as AlphaFold2.** When position b2b3.12 = I, dimer is predicted (A). When b2b3.12 = E, chemokine (referenced against Anc0) is predominantly predicted (B).

A.

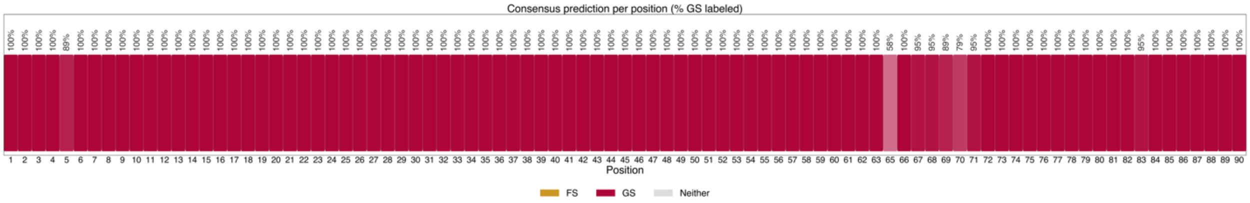


B.

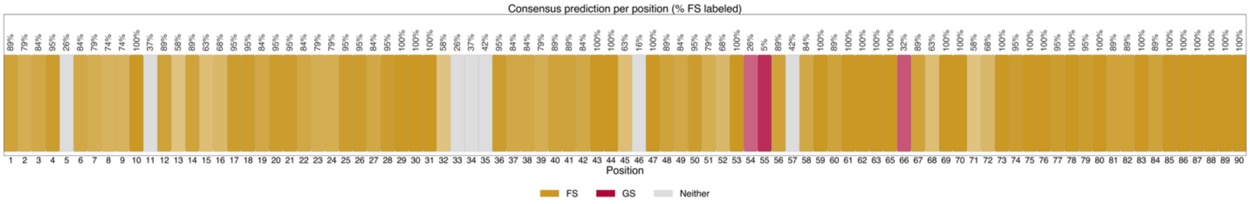


**Figure S10. AlphaFold3 exhibits similar gating behavior for KaiB as AlphaFold2, except L64E alone changes the conformation to FS rather than a conformation that differs from FS and GS.** When position 64 = L, GS is predicted (A). When 64 = E, FS is largely predicted (B).

**A.**

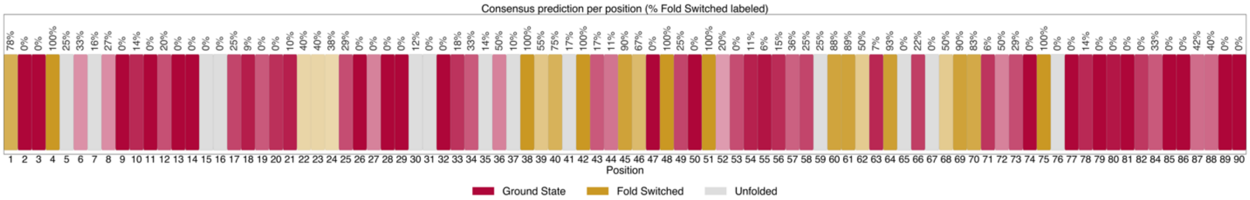


**B.**

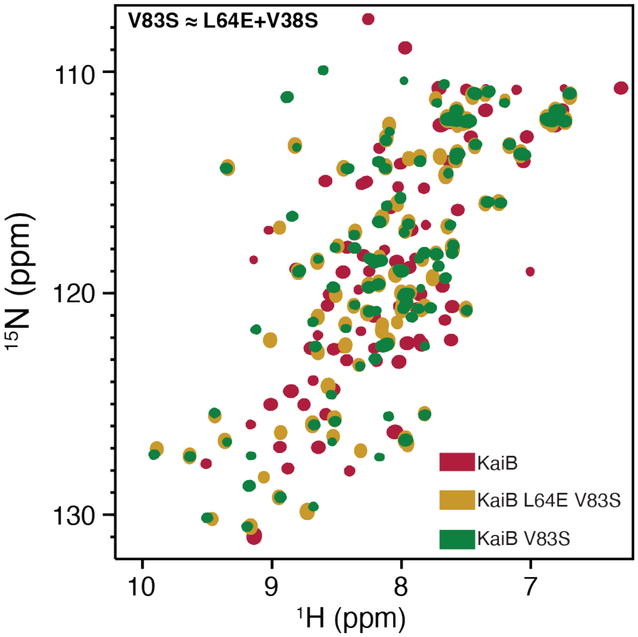


**Figure S11. The V83S mutation switches KaiB's major state from ground to fold-switched despite dominant ground state predictions.** (A). Strong gating behavior is not predicted for KaiB V83S as it is for KaiB L64E. Instead, most KaiB V83S variants are predicted ground state, rather than fold-switched or neither. (B). Nevertheless, substitution V83S shifts KaiB toward the fold-switched conformation assumed by KaiB L64E+V83S as evidenced by substantial peak overlap of these two variants. KaiB L64E V83S contours are enhanced to highlight overlap with V83S.

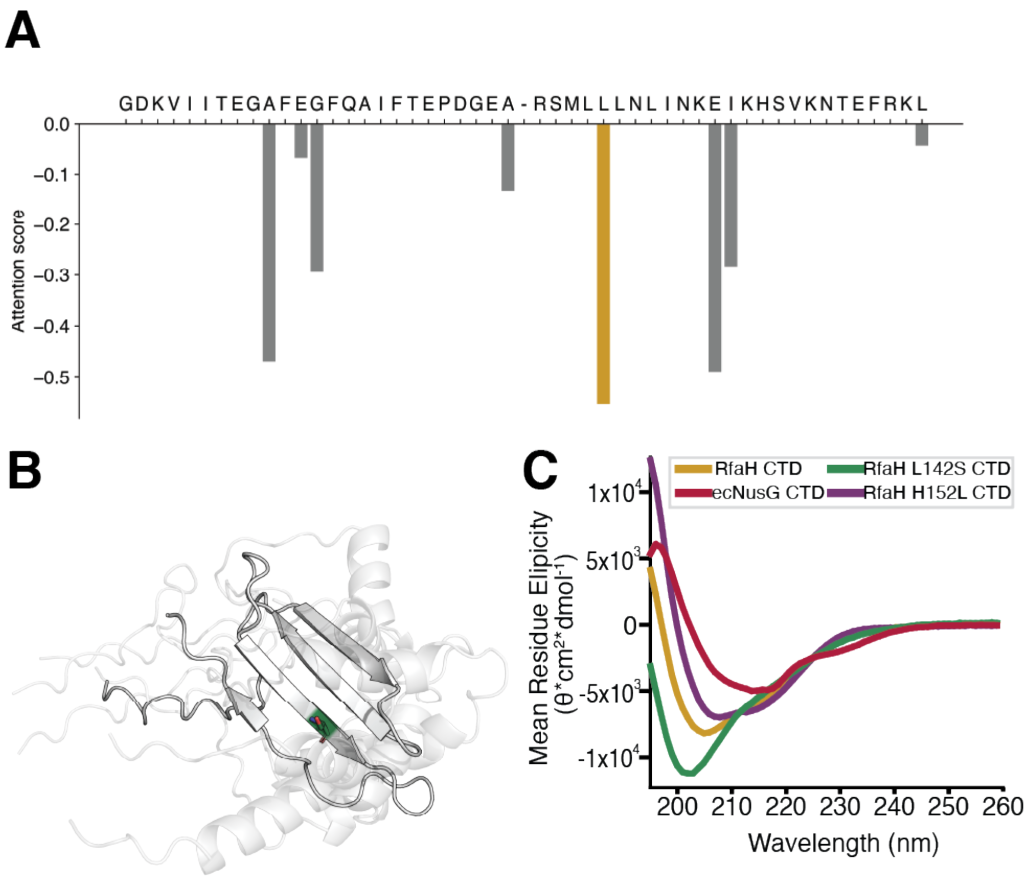


**Figure S12. CAAT identifies a mutation that gates RfaH CTD's folded state.** (A). CAAT identifies position 142 in RfaH CTD as the most strongly gating residue (yellow bar). (B). AlphaFold predicts heterogeneous, low-confidence structures in response to the L142S mutation to RfaH CTD. (C). Circular dichroism demonstrates that RfaH L142S is unfolded, in contrast to the unmutated RfaH, E. coli NusG, and RfaH H152L CTDs.

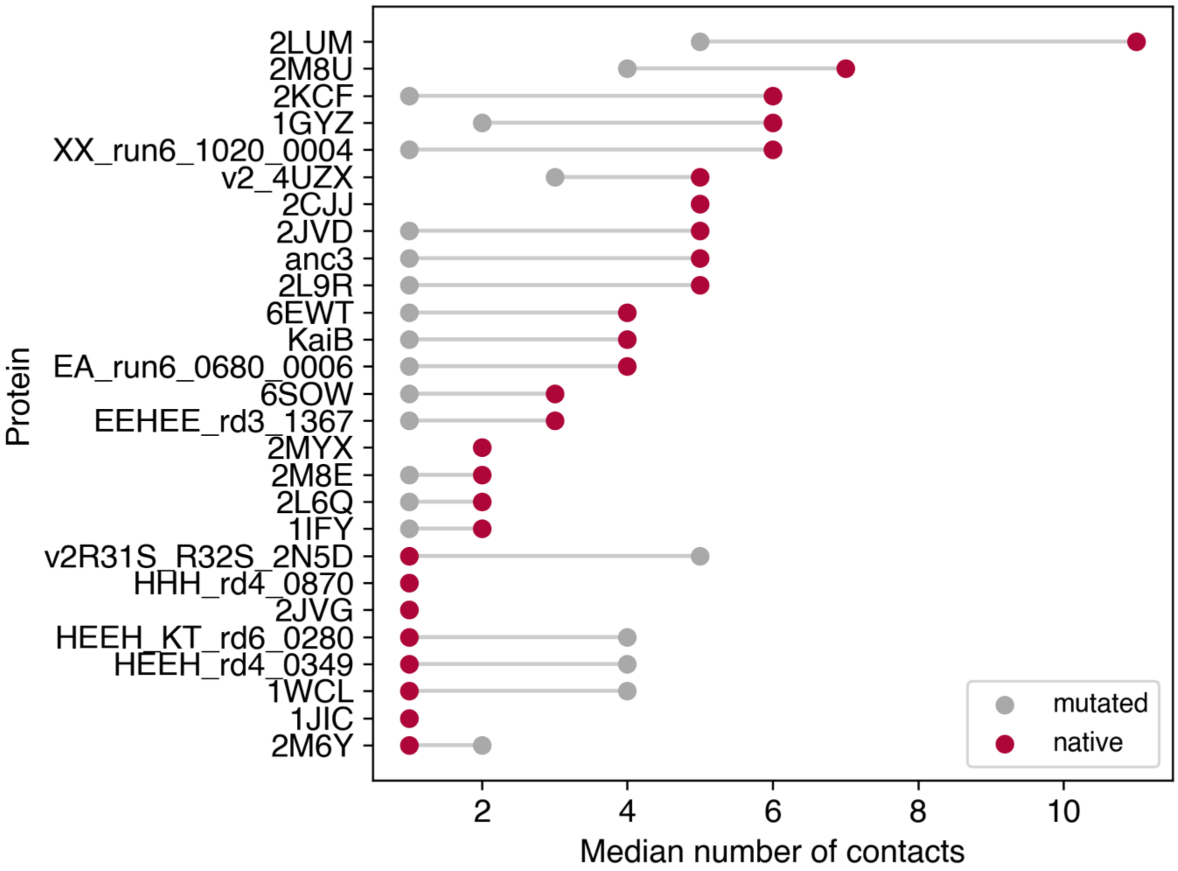


**Figure S13. A few proteins make new interaction networks rather than removing contacts.** These are v2R31S_R32S_2N5D, HEEH_KT_rd6_0280, HEEH_rd4_0349, 1WCL, and 2M6Y.

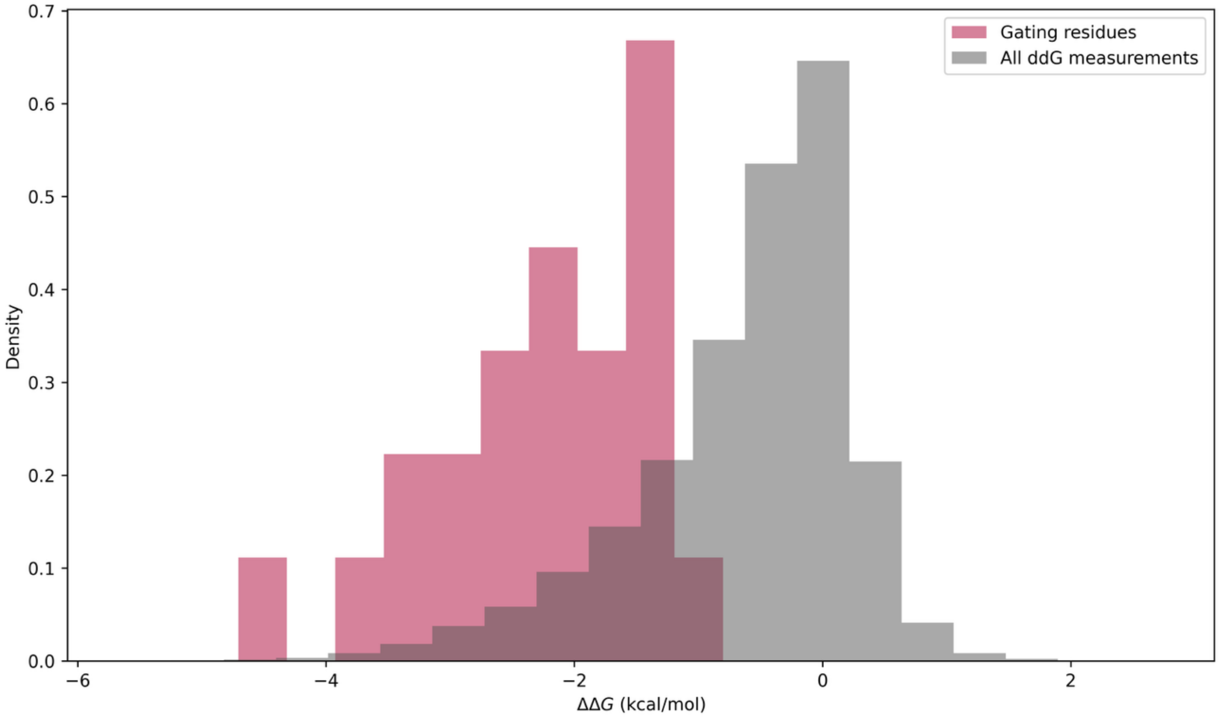


**Figure S14. Predicted gating residues destabilize proteins significantly more than by chance (KS test $p = 5 \times 10^{-14}$).**

A.

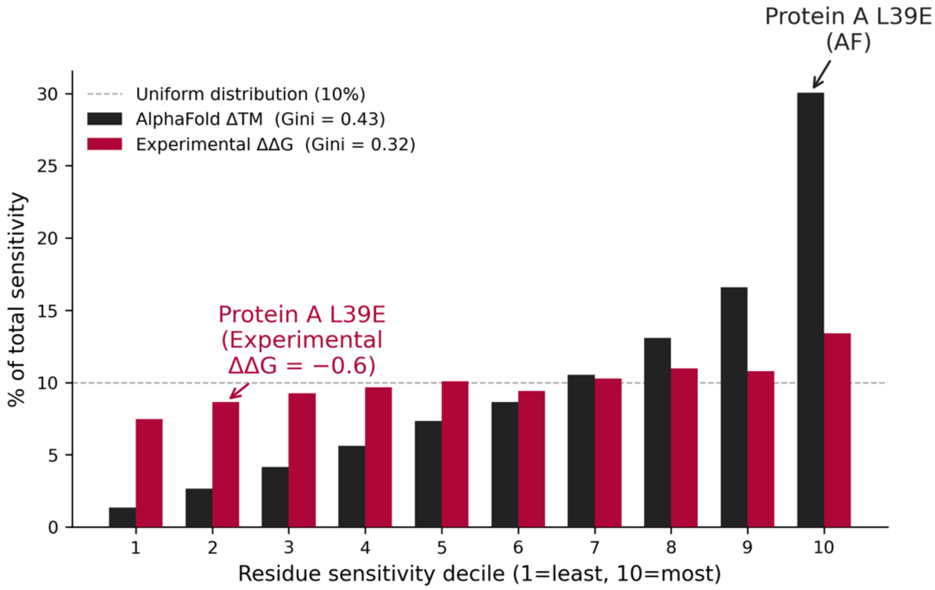


B.

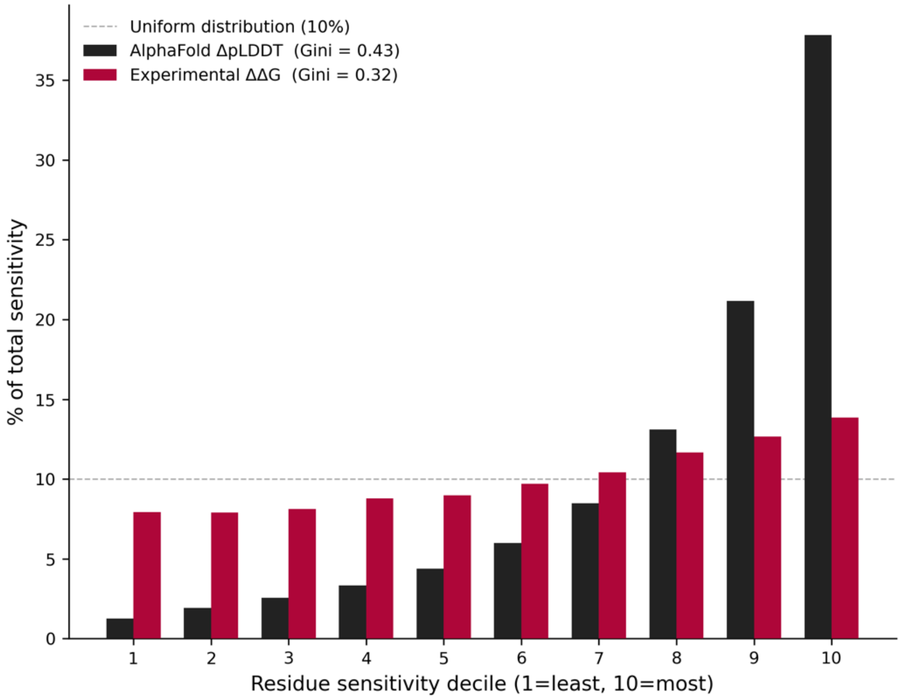


**Figure S15. AlphaFold sensitivity is more concentrated than experiment, and mutations predicted to be sensitive are not always so experimentally.** In both figures, residue sensitivity deciles are ranked by AlphaFold sensitivity for both prediction and experiment. (A) Is for sensitivity measured by change in TM-score. (B). Is for sensitivity measured by change in prediction confidence (pLDDT).

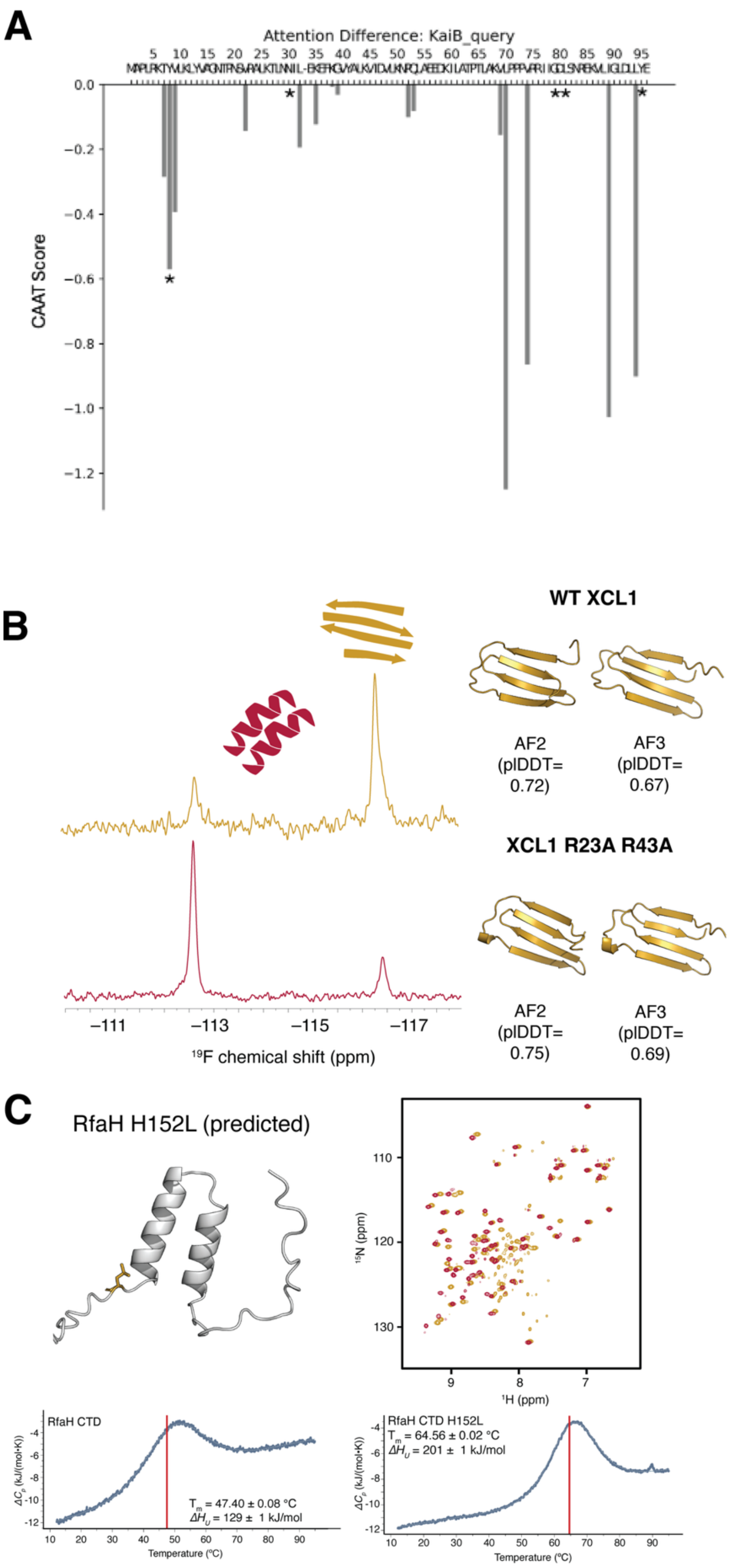


**Figure S16. AlphaFold can miss experimentally important mutations.** Caption overleaf.

**Figure S16** A. 5 mutations to *T. elongatus* KaiB switch its ground state to the fold-switched state (marked as asterisks). Nevertheless, AlphaFold predicts that these 5 mutations result in a ground-state structure. Four of the five sites have CAAT scores of 0 and the fifth has a moderate score. B. AlphaFold does not accurately predict effects of the XCL1 R23A R43A mutation. Though $^{19}$F NMR experiments indicate that the population shifts from largely dimer (wildtype) to largely chemokine (R23A R43A), both AlphaFold2 and AlphaFold3 predict the dimer fold for R23A R43A with higher confidence than wildtype. C. AlphaFold predicts a minor state for the RfaH H152L sequence, but the corresponding peak position diagrams shows the absence of the RfaH minor state for RfaH H152L (red) and stronger beta sheet peaks than for wildtype RfaH (yellow). Further, differential scanning calorimetry experiments show that the major beta-sheet state of RfaH CTD is substantially more stable in response to the H152L mutation than wildtype.

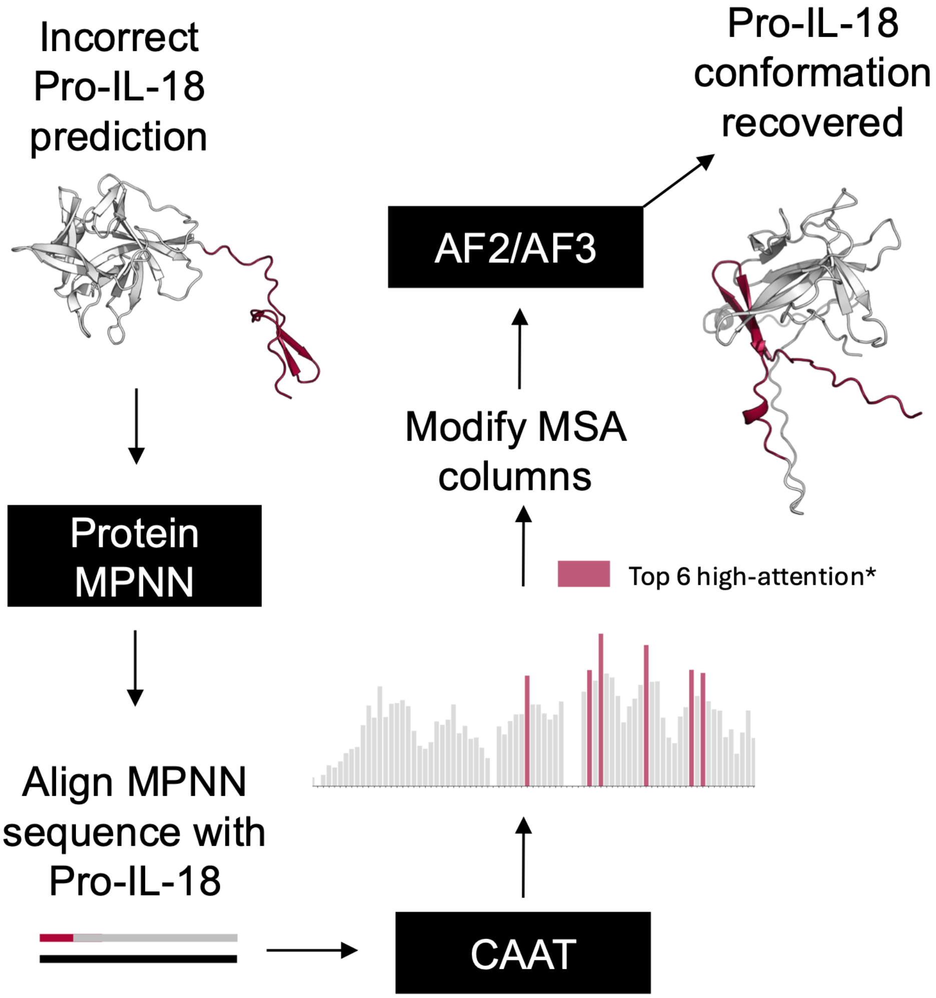


**Figure S17. CAAT-guided MSA editing recovers a pro-IL-18 conformation similar to experiment.** The incorrectly predicted pro-IL-18 conformation was passed into ProteinMPNN to generate a sequence (black) predicted to adopt the same incorrect fold. A two-sequence MSA containing pro-IL-18 as the target sequence (red) and the mature IL-18 sequence (gray) was then passed into CAAT to identify the most highly attended positions within the first 72 residues of pro-IL-18, which encompass the pro-domain insertion that distinguishes pro- from mature IL-18. Substitutions were chosen based on conservation between the pro-IL-18 and ProteinMPNN sequences. Four residues were conserved between the two sequences and hydrophobic (F38, V47, L56, F66); we reasoned that these residues anchor the incorrect mature conformation and therefore mutated the corresponding MSA columns — but not the target sequence itself — to charged residues (F38E, V47E, L56K, F66K). Two residues were not conserved between pro-IL-18 and the ProteinMPNN sequence; we reasoned that these may encode information favoring the correct pro-form and therefore mutated the MSA columns to match the pro-IL-18 identity (S46, P64). This combination of six MSA edits produced 15/15 AF2 predictions and the top-ranked AF3 prediction resembling the experimentally determined pro-IL-18 conformation.
